# Soil Bacterial and Fungal Response to Wildfires in the Canadian Boreal Forest Across a Burn Severity Gradient

**DOI:** 10.1101/512798

**Authors:** Thea Whitman, Ellen Whitman, Jamie Woolet, Mike D. Flannigan, Dan K. Thompson, Marc-André Parisien

## Abstract

Global fire regimes are changing, with increases in wildfire frequency and severity expected for many North American forests over the next 100 years. Fires can result in dramatic changes to C stocks and can restructure plant and microbial communities, with long-lasting effects on ecosystem functions. We investigated wildfire effects on soil microbial communities (bacteria and fungi) in an extreme fire season in the northwestern Canadian boreal forest, using field surveys, remote sensing, and high-throughput amplicon sequencing. We found that fire occurrence, along with vegetation community, moisture regime, pH, total carbon, and soil texture are all significant predictors of soil microbial community composition. Communities become increasingly dissimilar with increasingly severe burns, and the burn severity index (an index of the fractional area of consumed organic soils and exposed mineral soils) best predicted total bacterial community composition, while burned/unburned was the best predictor for fungi. Globally abundant taxa were identified as significant positive fire responders, including the bacteria *Massilia sp*. (64× more abundant with fire) and *Arthrobacter sp*. (35×), and the fungi *Penicillium sp*. (22×) and *Fusicladium sp.* (12×). Bacterial and fungal co-occurrence network modules were characterized by fire responsiveness as well as pH and moisture regime. Building on the efforts of previous studies, our results identify specific fire-responsive microbial taxa and suggest that accounting for burn severity improves our understanding of their response to fires, with potentially important implications for ecosystem function.

## Introduction

The boreal forests of Canada hold roughly 10% (between 168–200 Pg) of total global terrestrial carbon (C) stocks [1] and cover about 55% of the country’s landmass [2]. Wildfire is a natural part of these ecosystems, playing a key role in structuring vegetation communities and affecting above- and belowground C stocks through combustion and persistent effects on the subsequent forest recovery [3–6]. Much of this soil C is stored in peatlands (peat-forming wetlands), which cover as much as 50% of the land surface in some boreal forest landscapes [7], and act as substantial sources of methane [8]. Because soil microbes play an important role in these ecosystems, both governing soil C cycling and also interacting directly with plants, it is important to understand how soil microbes are affected by wildfire.

Microbial response to fire has been studied for over a century [9–10]. Across studies, wildfires usually decrease soil microbial biomass [11–13], and microbial communities can take decades to recover to pre-fire states [13–15]. There are numerous mechanisms through which fires can affect soil microorganisms. Briefly, we can organize these mechanisms into three categories: (1) fire directly killing microbes and destroying microbial habitat; (2) altered post-fire chemicophysical environment (*e.g.*, increased pH, changes to water permeability, changed nutrient inputs or competition for nutrients from plants); (3) altered post-fire biological environment (*e.g.*, competitors removed, differential survival of taxa, or loss of symbiont plants) [16–17]. Investigating specific fire-responsive taxa and beginning to decipher their ecological strategies may help us predict the long-term ecological and biogeochemical effects of fire.

Although numerous field studies have investigated how microbes are affected by fires, few consider multiple ecosystem types, more than one fire, or how these effects change with increasing burn severity. Wildfires can range from lightly burned surface fires with no tree mortality, to fires that kill all trees and remove all or most of the soil litter layer (O horizon). As fire regimes around the world are affected by climate change [4, 18], fire frequency, size, and intensity (and thus, indirectly, burn severity) are expected to increase in many areas of the boreal forest [18–19]. Burn severity is the degree of fire-induced change to vegetation and soils [20, 21]. When past studies have considered burn severity, some have observed that increasing burn severity is associated with reduced fungal abundance [22] or changes to community composition [23]. For bacteria, while some studies have found only negligible effects of burn severity on community composition [24] others have noted distinct changes to bacterial community composition with different levels of burn severity [25]. These changes to microbial communities could have interesting interactive effects with plant colonization post-fire. For example, Knelman *et al*. [26] reported interactive effects between burn severity (low *vs*. high) and the effect of plant colonization (*Corydalis aurea* presence *vs*. absence) on bacterial communities and potential activity in a Colorado, USA *Pinus ponderosa* forest.

There are numerous metrics of burn severity used by fire scientists, each of which represents different aspects of the effects of fire. Field-based ground metrics include single measurements such as percent exposed mineral soil or mean duff depth (O horizon); the burn severity index (BSI) integrates the burn severity of the forest floor and soil surface [27]. Canopy fire severity index (CFSI) estimates the intensity of the combustion of large trees [28]. Composite burn index (CBI) is a generalized measure of burn severity, mortality, and combustion across all forest strata from soils to large trees [29–30]. The remotely-sensed relativized burn ratio (RBR [21]) combines satellite imagery from before and after burns, capturing changes in reflectance due to vegetation combustion and mortality, and combustion of organic soil and changes in soil moisture. Few studies have investigated the relative utility of these different burn severity metrics for predicting microbial community response to fire. Determining the effects of fire on soil microbial communities across a wide range of sites and assessing the utility of different burn severity metrics could underpin efforts to predict and characterize the effects of wildfires and changing wildfire regimes on soil microbes in boreal upland and wetland soils.

The first objective of our study was to determine the relative importance of soil, vegetation, and wildfire severity metrics in predicting soil microbial community composition one year post-fire across five vegetation types in the boreal forests of northwestern Canada. Our studied regions have high pedodiversity, spanning wide ranges of pH, texture, and organic horizon thicknesses. We hypothesized that vegetation community and soil pH would be the strongest predictors of microbial community composition, while the effect of fire might not be a significant factor after controlling for vegetation community and soil properties across such a wide range of sites. If fire were to be found to be a significant predictor, we hypothesized that burn metrics associated with the ground surface would outperform remotely sensed metrics or canopy burn metrics as predictors of microbial community composition. The second objective of our study was to identify specific fire-responsive taxa. To achieve these objectives, we sampled six large wildfires one year after they burned in the Northwest Territories and northern Alberta, Canada, characterizing soil, vegetation, and fire properties, and sequencing microbial (bacterial/archaeal and fungal) communities using the ribosomal RNA gene.

## Methods

### Study region

We selected sites in the Northwest Territories and northern Alberta (Wood Buffalo National Park), Canada, and sampled them one year post-fire, in 2015 (Figure 1; Supplemental Table 1; Supplemental Note 1). The fires and the drivers of burn severity are described in detail in Whitman *et al*. [31], while their effects on understory vegetation are described in detail in Whitman *et al*. [32]. The study region has long, cold winters and short, hot summers, with mean annual temperatures between −4.3 °C and −1.8 °C and annual precipitation ranging from 300 to 360 mm [33–34]. The fire regime of the study region includes infrequent stand-replacing fires every 40-350 years on average [35]. The majority (~97%) of the burned area is contributed by ~3% of fires [36]. The six large wildfires in this study ranged in size from 14 000 to 700 000 ha. The soils in these regions are mostly classified as Typic Mesisols (32 sites), Orthic Gleysols (16 sites), or Orthic Gray Luvisols (8 sites) (Soil Landscapes of Canada map v.3.2). The sampled sites span a wide range of soil properties, with pH values ranging from 3.2 (wetlands) to 8.1 (uplands with calcareous plant material), total C ranging from 0.5% (mineral horizon) to 52% (organic horizon), and a wide range of textures (Supplemental Table 2). We classified vegetation communities for each upland site as being jack pine-dominated (*Pinus banskiana* Lamb.), black spruce-dominated (*Picea mariana* (Mill.)), or composed of a mix of coniferous and broadleaf trees (“mixedwood”). We classified vegetation communities for wetlands as open or treed [37].

**Figure 1.**
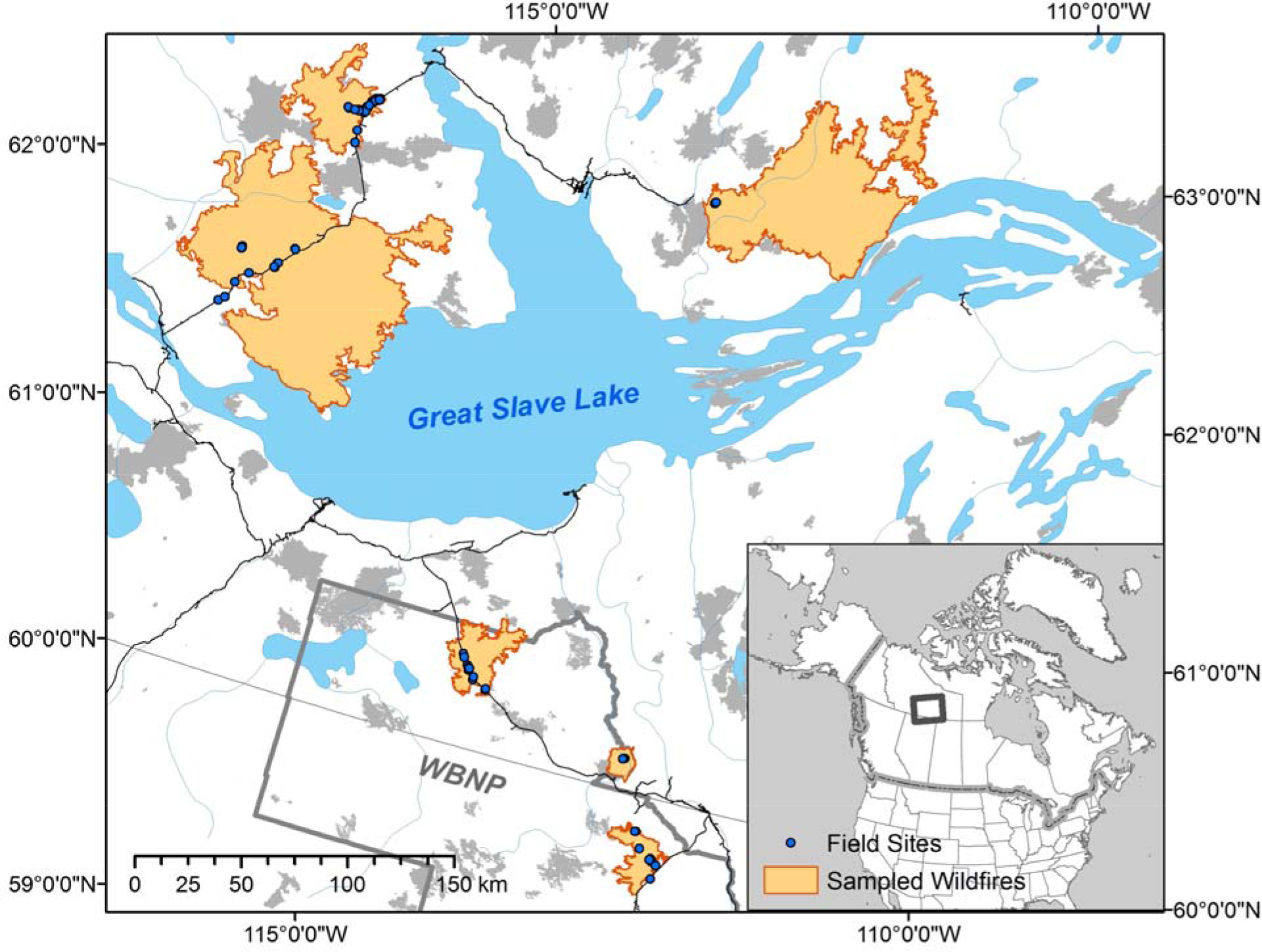
Study region of northern Alberta and the Northwest Territories, Canada, including Wood Buffalo National Park (WBNP – grey outline). Blue points indicate sampled sites, orange shapes indicate sampled fires, and grey shapes indicate other 2014 fires in the region. Inset indicates relative location within North America.

### Site assessment methodologies

Sites were selected and characterized as described in detail by Whitman *et al*. [31–32]. Briefly, field sites were selected to represent the local range of burn severity and vegetation communities, resulting in a total of 50 burned field sites. We selected an additional 12 control sites (not burned within the last 38 years before sampling, mean time since fire 95 years; “unburned”), chosen to reflect the range of vegetation communities sampled in the burned plots, for a total of 62 sites. At each site we established a 30 × 30 m square plot with 10 × 10 m subplots at the four corners. We measured post-fire organic horizon depth (up to 10 cm) at the inner corners of the 10 × 10 m subplots. Understory vegetation percent cover was assessed in five 1 × 1 m plots at the same four points as organic soil depth, and at the plot centre [32]. We assessed burn severity in the four subplots (described in detail in [31]; Supplemental Table 1), using severity metrics of canopy fire severity index (CFSI [28]), burn severity index (BSI [27]), and percent exposed mineral soil. We also assessed the composite burn index (CBI; understory, overstory, and mean [29–30]) in the entire 30 × 30 m plot area. We used the relativized burn ratio (RBR) to represent remotely sensed burned severity at each site [21]. RBR was produced using multispectral Landsat 8 Operational Land Imager and Landsat 5 Thematic Mapper images (Landsat Level-1 imagery, courtesy of the USGS).

At each plot, we took soil cores (5.5 cm diameter, 13.5 cm depth) at three locations (centre, SW and NE subplots). Soil cores were gently extruded and separated into organic (O) horizons (where present) and mineral (M) horizons (where present in the top 13.5 cm of soil profile). The three samples were pooled by horizon at each site and mixed gently by hand in a bag. From these site-level samples, sub-samples were collected for microbial community analysis and stored in LifeGuard Soil Preservation solution (QIAGEN, Germantown, MD) in a 5 mL tube (Eppendorf, Hamburg, Germany). Tubes were kept as cold as possible while in the field (usually for less than 8 h, but up to 2 days for remote sites) and then stored frozen. The remaining soil samples were air-dried and analyzed for a range of properties, including, pH and total C (Supplemental Table 2; Supplemental Note 1).

### DNA extraction, amplification, and sequencing

Duplicate DNA extractions were performed for each sample, with two blank extractions for every 24 samples (half of which were sequenced), using a DNEasy PowerLyzer PowerSoil DNA extraction kit (QIAGEN, Germantown, MD) following manufacturer’s instructions. Extracted DNA was amplified in triplicate PCR, targeting the 16S rRNA gene v4 region (henceforth, “16S”) with 515f and 806r primers [38], and targeting the ITS2 gene region with 5.8S-Fun and ITS4-Fun primers [39] with barcodes and Illumina sequencing adapters added as per [40] (all primers in Supplemental Tables 3-5). The PCR amplicon triplicates were pooled, purified and normalized using a SequalPrep Normalization Plate (96) Kit (ThermoFisher Scientific, Waltham, MA). Samples, including blanks, were pooled and library cleanup was performed using a Wizard SV Gel and PCR Clean-Up System A9282 (Promega, Madison, WI). The pooled library was submitted to the UW Madison Biotechnology Center (UW-Madison, WI) for 2×250 paired end (PE) Illumina MiSeq sequencing for the 16S amplicons and 2×300 PE for the ITS2 amplicons. (See Supplemental Note 1 for full details.)

### Sequence data processing and taxonomic assignments

For 16S reads, we quality-filtered and trimmed, dereplicated, learned errors, determined operational taxonomic units (OTUs), and removed chimeras using dada2 [41] as implemented in R. For ITS2 reads, we first merged reads using PEAR [42], and then performed the same steps as for 16S. These sequence processing steps were performed on the UW-Madison Centre for High Throughput Computing cluster (Madison, WI). After confirming that the paired DNA extraction replicates were very similar to each other, we combined the community composition data from paired extractions additively and proceeded with a single sequencing dataset for each soil sample. Taxonomy was assigned to the 16S reads using a QIIME2 [43] scikit-learn feature classifier trained on the 515f-806r region of the 99% ID OTUs from the Silva 119 database [44]. We used BLAST to determine the closest > 97% ID match, if present, in the database of globally abundant bacterial phylotypes [45]. For the ITS2 reads, we first ran them through ITSx [46] to identify fungi and to remove plant sequences, and then assigned taxonomy using the UNITE species hypothesis 99% threshold database version 7.2 [47], using the parallel_assign_taxonomy_uclust.py script in QIIME1 [43] with default settings to the genus level. We also classified ITS2 taxonomic assignments using the FunGuild database [48]. (See Supplemental Note 1 for full details of bioinformatics, including sequences retained at each step.)

### Quantitative PCR

To estimate the relative abundance of bacteria *vs.* fungi in a given sample, extracted DNA was amplified via quantitative PCR (qPCR) in triplicate, targeting the 16S rRNA gene v4 region with 515f and 806r primers [49] and targeting the 18S gene region with FR1 and FF390 primers [50] (Supplemental Note 1; qPCR primers in Supplemental Table 3; raw Cq values and calibration curves in Supplemental File 1).

### Bioinformatics and statistics

We worked primarily in Jupyter notebooks, with phyloseq [51], ggplot [52], and dplyr [53] being instrumental in working with the data in R [54] (See Supplemental Note 1 for full bioinformatics details).

We compared community composition across samples using Bray-Curtis dissimilarities on Hellinger-transformed relative abundances [55], which we represented using NMDS ordinations. We tested for significant effects of vegetation community, moisture regime (as a continuous variable), pH, total C, texture (% sand), and burned/unburned using a permutational multivariate ANOVA (PERMANOVA; the adonis function in vegan [56]). Because the order of the terms in the PERMANOVA model affects the partial R^2^ of a given term, to compare the relative explanatory power of each component, we also compared the R^2^ of single-component models for each factor. We predicted 16S rRNA gene copy numbers using the ribosomal RNA operon database (rrnDB) [57]. We calculated the abundance-weighted mean predicted copy number for each sample using the approach of Nemergut *et al*. [58]. We tested for the relationship between weighted mean predicted copy number for each sample and burn severity (understory composite burn index) with a linear model. To compare our findings with those of Holden *et al*.[23], we calculated the mean understory CBI value for all sites at which each OTU within that phylum was present, and determined whether there were significant differences in these values between different fungal phyla, using an ANOVA and Tukey’s HSD for multiple comparison correction. We tested whether vegetation community dissimilarity was significantly correlated with bacterial or fungal community dissimilarity for all pairs of sites in mineral and in organic soil horizons using Mantel tests, with 999 permutations.

To examine the relative explanatory power of different burn severity metrics for predicting microbial community composition, we used a simple linear model, which controlled for parameters we expected to influence community composition – vegetation community, moisture regime, pH, texture (% sand), and total C – and then tested the inclusion of each severity metric, comparing the partial R^2^ values for the severity metric. The severity metrics we tested included: Burned/Unburned, RBR, CFSI, CBI, Understory CBI, Overstory CBI, BSI, % exposed mineral soil, and mean duff depth. To determine whether communities become increasingly dissimilar with increasing burn severity, we fit a linear model to Bray-Curtis dissimilarity (to unburned samples from sites with the same vegetation community and soil horizon) *vs*. BSI. We calculated the relationship between BSI and log(16S abundance : 18S abundance) using a linear model in R. We also did the same with pH and log(16S abundance : 18S abundance).

We estimated richness and its associated standard error in each sample using the *breakaway* function in R [59]. We determined which OTUs were significantly enriched in burned plots (*vs.* unburned plots) using metagenomeSeq [60], after controlling for (including as variables) vegetation community (categorical variable), pH (continuous variable), and %C (continuous variable), resulting in an estimate of the log_2_-fold change in the abundance of each OTU in burned vs. unburned plots, across samples. For a small subset of OTUs, we investigated the relationship between their log(relative abundance) and BSI using a linear model.

To determine which fungal and bacterial OTUs and understory vegetation co-occurred across samples, we used a network analysis approach, following Connor *et al*. [61] to avoid false positives and establish conservative network cutoff parameters. After simulating a null model network to choose an appropriate rho value, we determined a consensus network by adding random tie-breaking noise to the matrix 2000 times, selecting only the co-occurrences that occurred in 95% of the 2000 replications. We determined standard network characterization metrics [62–66], including modularity using random walks, and plotted the network using igraph R package [67].

## Results

### Community-level characteristics and predictors

Bacterial and fungal communities were dominated by typical soil organisms [45, 68] (Supplemental Figures 1-4). All tested factors (vegetation community, moisture regime, pH, total C, texture, and burned/unburned) were significant predictors of community composition for both bacteria (Figure 2 and Supplemental Figure 5) and fungi (Figure 2 and Supplemental Figure 6) in the combined model (PERMANOVA, p<0.015 for all factors). Moisture regime and vegetation community provided the most explanatory power for bacteria and for fungi (R^2^ values between 0.12 and 0.15 for the individual models). After these factors, for bacteria, pH provided the most explanatory power (R^2^_pH_ = 0.07; Figure 2A). For fungi, carbon provided the most explanatory power (R^2^_C_ = 0.05; Figure 2B).

**Figure 2.**
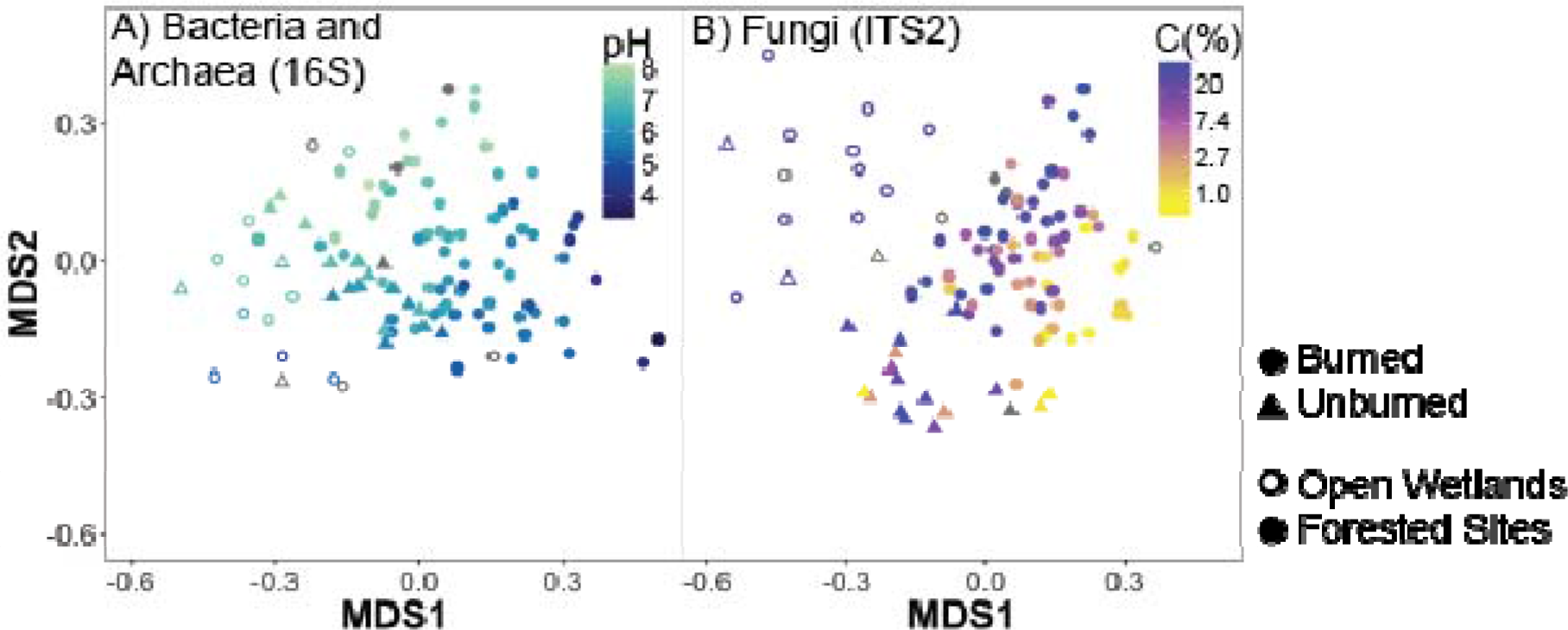
(A) NMDS ordination of Bray-Curtis distances between bacterial/archaeal (16S rRNA gene v4 region) communities for all samples (k=2, stress=0.16). Circles indicate burned plots, while triangles indicate unburned plots, and open points indicate open wetland sites. Points are shaded by pH, with darker colours indicating lower pH values. Grey points indicate samples for which pH values were not attainable, due to insufficient sample mass. (B) NMDS ordination of Bray-Curtis distances between fungal (ITS2) communities for all samples (k=3, stress=0.14). Circles indicate samples from burned plots, whereas triangles indicate unburned plots, and open points indicate open wetland sites. Points are shaded by C content, with darker colours indicating higher C. Note logged colour scale. Grey points indicate samples for which C values were not attainable, due to insufficient sample mass.

Different fungal phyla were present at different levels of burn severity. OTUs within *Chytridiomycota* and *Mucoromycota* occurred at sites with significantly higher mean BSI than *Ascomycota, Basidiomycota*, or *Rozellomycota* in upland sites (Supplemental Figure 7A). In wetland sites, OTUs within *Basidiomycota* occurred at sites with significantly lower mean BSI than for all other phyla except *Rozellomycota*. This trend largely mirrored that of the same approach with moisture regime instead of BSI (Supplemental Figure 7B), but after controlling for moisture regime, similar trends persisted (Supplemental Figure 8).

There was a significant positive relationship between burn severity and weighted mean predicted 16S copy number (Figure 3A). The OTUs identified as positive fire-responders had significantly higher mean predicted 16S copy number than those identified as negative fire-responders (3.6 *vs*. 2.6, p=0.01).

Bacterial and fungal communities become increasingly dissimilar from unburned sites with increasing burn severity in upland sites (p<0.001, Figure 3B and 3C). We did not detect such a relationship for wetland sites (p>0.05) (Supplemental Figure 9). Across all sample pairs, there was a significant positive relationship between understory vegetation community dissimilarity and organic horizon microbial community dissimilarity in wetlands but not in uplands (Supplemental Figure 10) - *i.e.*, wetland sites with similar understory plant communities have similar microbial communities.

**Figure 3.**
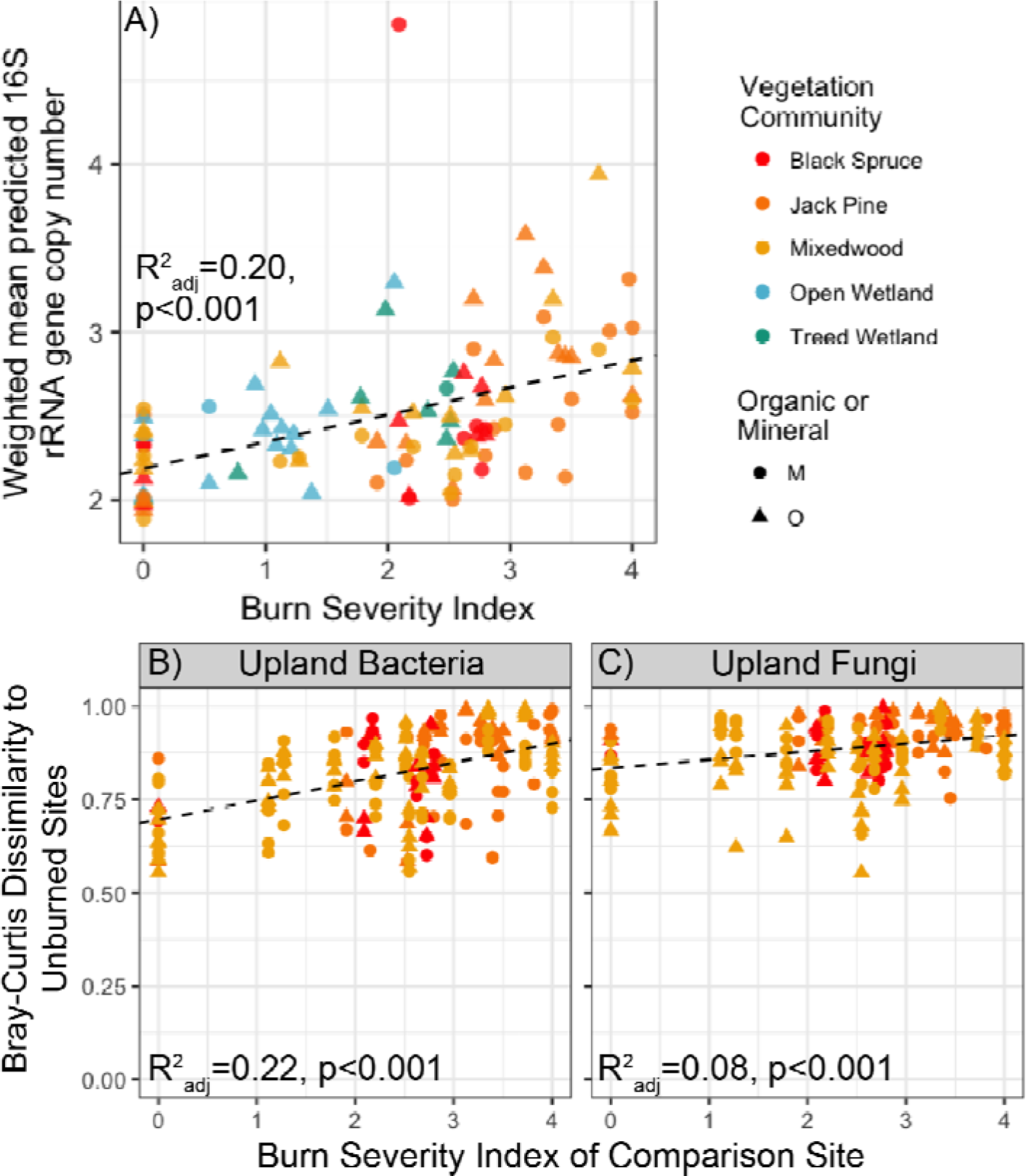
(A) Relationship between weighted mean predicted 16S rRNA gene copy number and burn severity index (BSI). Dashed line indicates linear fit (y = 2.19+0.16x, R^2^_adj_ = 0.20, p<0.001). (B) Bray-Curtis dissimilarity to unburned sites (within the same vegetation community and the same soil horizon type) for bacteria in uplands vs. burn severity index of comparison sites. Dashed lines indicate linear fit (y = 0. 0.05 × + 0.70, p<0.001, R^2^_adj_ = 0.22). (C) Bray-Curtis dissimilarity to unburned sites (within the same vegetation community and the same soil horizon type) for fungi in uplands vs. burn severity index of comparison sites. Dashed lines indicate linear regressions (y = 0.02 × + 0.83, p<0.001, R^2^_adj_ = 0.08). For all figures, points are coloured by vegetation community; circles represent mineral horizon samples, triangles represent organic horizon samples. Equivalent figures for wetlands for (B) and (C) are found in Supplemental Figure 9.

**Table 1.** Predictive value of burn severity metrics in models of Hellinger-transformed microbial community Bray-Curtis dissimilarities, after controlling for vegetation community, moisture regime, pH, total C, and texture (% sand). The best models for each group are highlighted in bold text. N_bacteria_ = 94, N_fungi_ = 92, except for Overstory CBI, where N_bacteria_ = 90, N_fungi_ = 88

| Severity metric | $p_{\text{severity}}$ | Partial $R^2_{\text{severity}}$ | $R^2_{\text{full}}$ |
| --- | --- | --- | --- |
| *Bacteria* (16S rRNA gene v4 region) |  |  |  |
| **Burned/Unburned** | **0.001** | **0.037** | **0.29** |
| Relativised Burn Ratio | 0.001 | 0.028 | 0.29 |
| Canopy Fire Severity Index | 0.001 | 0.020 | 0.28 |
| Composite Burn Index (CBI) | 0.001 | 0.032 | 0.29 |
| Understory CBI | 0.001 | 0.033 | 0.29 |
| Overstory CBI | 0.001 | 0.031 | 0.29 |
| **Burn Severity Index** | **0.001** | **0.039** | **0.30** |
| % exposed mineral soil | 0.001 | 0.026 | 0.28 |
| Mean duff depth | 0.034 | 0.013 | 0.27 |
| *Fungi* (ITS2) |  |  |  |
| **Burned/Unburned** | **0.001** | **0.044** | **0.28** |
| Relativised Burn Ratio | 0.001 | 0.036 | 0.27 |
| Canopy Fire Severity Index | 0.001 | 0.030 | 0.27 |
| Composite Burn Index (CBI) | 0.001 | 0.040 | 0.28 |
| Understory CBI | 0.001 | 0.041 | 0.28 |
| Overstory CBI | 0.001 | 0.039 | 0.27 |
| **Burn Severity Index** | **0.001** | **0.043** | **0.28** |
| % exposed mineral soil | 0.001 | 0.026 | 0.26 |
| Mean duff depth | 0.001 | 0.021 | 0.26 |

All burn severity metrics added significant (but relatively little) additional predictive power to the model explaining microbial community composition. For bacteria, burn severity index was the best predictor of microbial community composition (partial R^2^_BSI_ = 0.040, p=0.001), marginally better than burned/unburned (Table 1). For fungi, a simple burned/unburned metric was the best predictor of microbial community composition (partial R^2^_B/U_ = 0.045, p=0.001), marginally better than burn severity index (Table 1).

There was not a significant relationship (p>0.05) between BSI (or other measures of burn severity) and 16S:18S gene copy numbers as determined by qPCR (Supplemental Figure 11). However, there was a significant (p<0.001) but weak positive relationship between pH and 16S:18S gene copy numbers as determined by qPCR, suggesting an increasing relative abundance of fungi in more acidic soils (Supplemental Figure 12).

There were no significant detectable changes in estimated richness across different vegetation communities or with increasing burn severity for bacteria (Supplemental Figure 13) or fungi (Supplemental Figure 14).

### Specific fire-responsive microbes

There were wide ranges of responses to wildfire within individual phyla. Numerous bacterial OTUs were identified as being significantly enriched (160 OTUs) or depleted (133 OTUs) in burned *vs*. unburned sites, after controlling for vegetation community, total C, and pH (Figure 4A; Supplemental Table 6; Supplemental Figures 15 and 16). About half of the responsive OTUs were at least 97% ID similar to the globally dominant phylotypes identified by Delgado-Baquerizo *et al*. [45] (Supplemental Figures 17 and 18). The most abundant bacterial OTU across samples (average 4% in burned samples *vs*. average 0.09% in unburned samples) was identified as a positive fire responder and was classified as an *Arthrobacter sp*. The third most abundant OTU (average 2% of the community in burned samples and not detected in unburned samples) was also identified as a positive fire responder and was classified as a *Massilia sp*. Of the bacterial taxa identified as being fire-responsive, different OTUs also showed different trends with burn severity (Figure 4B): the relative abundance of the *Arthrobacter sp.* (OTU sq1 and sq7 (we use the arbitrarily numbered “sq#” to distinguish specific OTUs)) increased with increasing burn severity (p<0.005). *Aeromicrobium* (OTU sq8), *Blastococcus* (OTU sq20), and *Massilia* (OTU sq3) also increased in relative abundance with increasing burn severity (p<0.05), with the *Massilia* and *Blastococcus* OTUs not even being detectable at any unburned sites.

**Figure 4.**
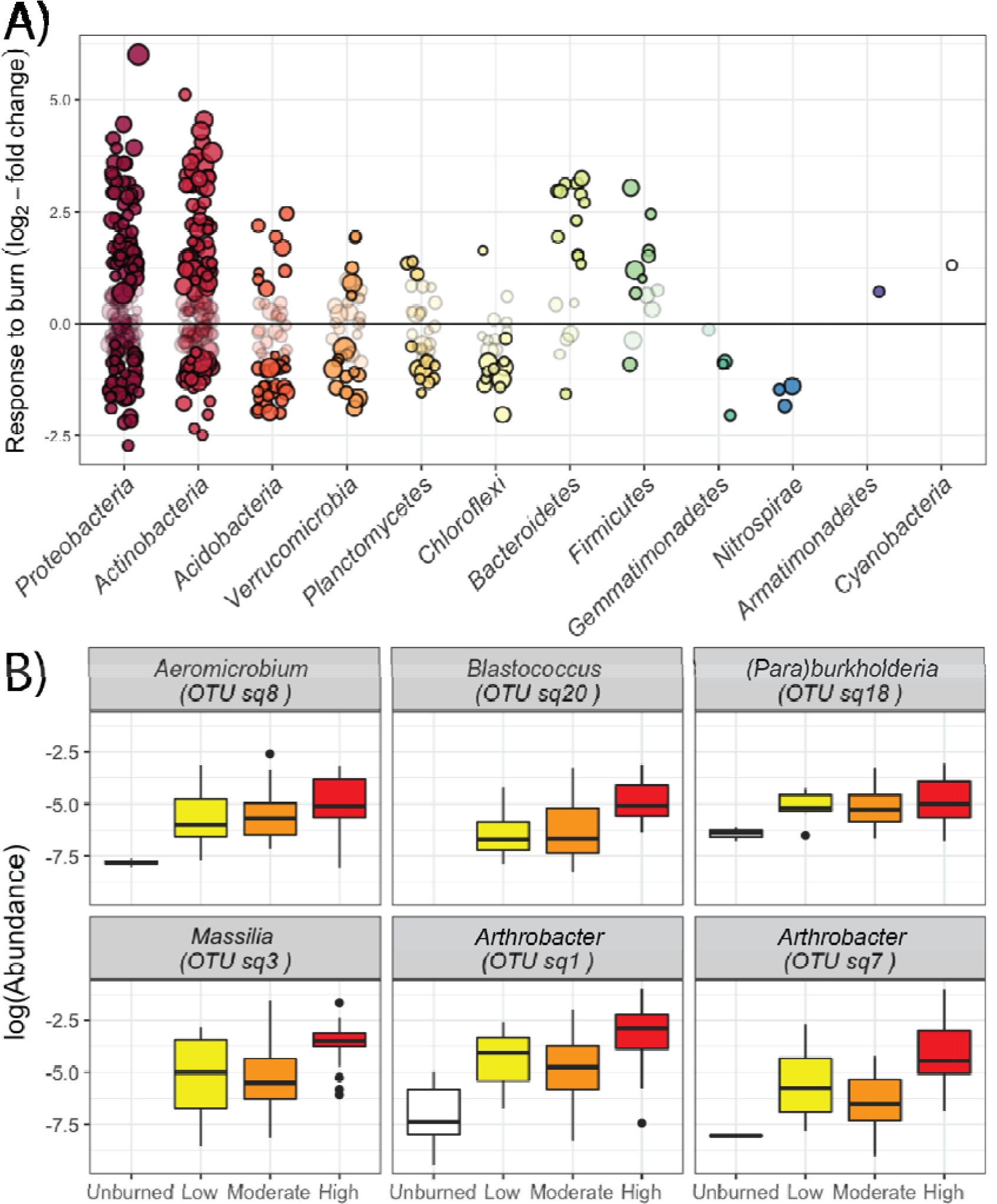
Bacterial response to fire. (A) Log_2_-fold change in burned *vs*. unburned plots, controlling for vegetation community, total C, and pH. Each point represents a single 16S rRNA gene v4 region OTU, and the size of each point represents the mean relative abundance of that OTU across all samples. Faint points represent OTUs that were not significantly different in abundance in burned *vs*. unburned plots. (B) Relative abundance (note log scale) of selected fire-responsive bacterial OTUs across BSI ranges (unburned=0, 0-2 low, 2-3 moderate, 3-4 high).

Numerous fungal OTUs were identified as being significantly enriched (79 OTUs) or depleted (60 OTUs) in burned *vs*. unburned sites, after controlling for vegetation community, total C, and pH (Figure 5A; Supplemental Table 7 and Supplemental Figure 19). There were wide ranges of responses within classes, with the exceptions of *Dothideomycetes* and *Cystobasidiomycetes* OTUs (which tended to be enriched in burned sites) and *Mortierellomycotina* subdivision (which tended to be depleted within burned sites). Certain fungal OTUs also stood out as fire responders. For example, fire-responsive OTUs included *Neurospora* and *Geopyxis* – genera that include well-known fire-responsive fungi. The third most abundant OTU was identified as a fire responder and was classified as *Penicillium sp*. Notable negative fire responders included three *Oidiodendron* OTUs (Supplemental Figure 20). Different fire-responsive OTUs showed different trends with burn severity (Figure 5B): the relative abundance of *Penicillium* (OTUs sq4 and sq65) and one *Fusicladium* (OTU sq24) increased significantly with increasing burn severity (p<0.05), whereas another *Fusicladium*, *Calyptrozyma*, and *Sordariomycetes* had a significant fire response, but did not continue to increase in relative abundance with increasing fire severity (p=0.11, 0.15, and 0.23, respectively). There were not consistent response patterns within putative mycorrhizal fungi (Supplemental Table 7).

**Figure 5.**
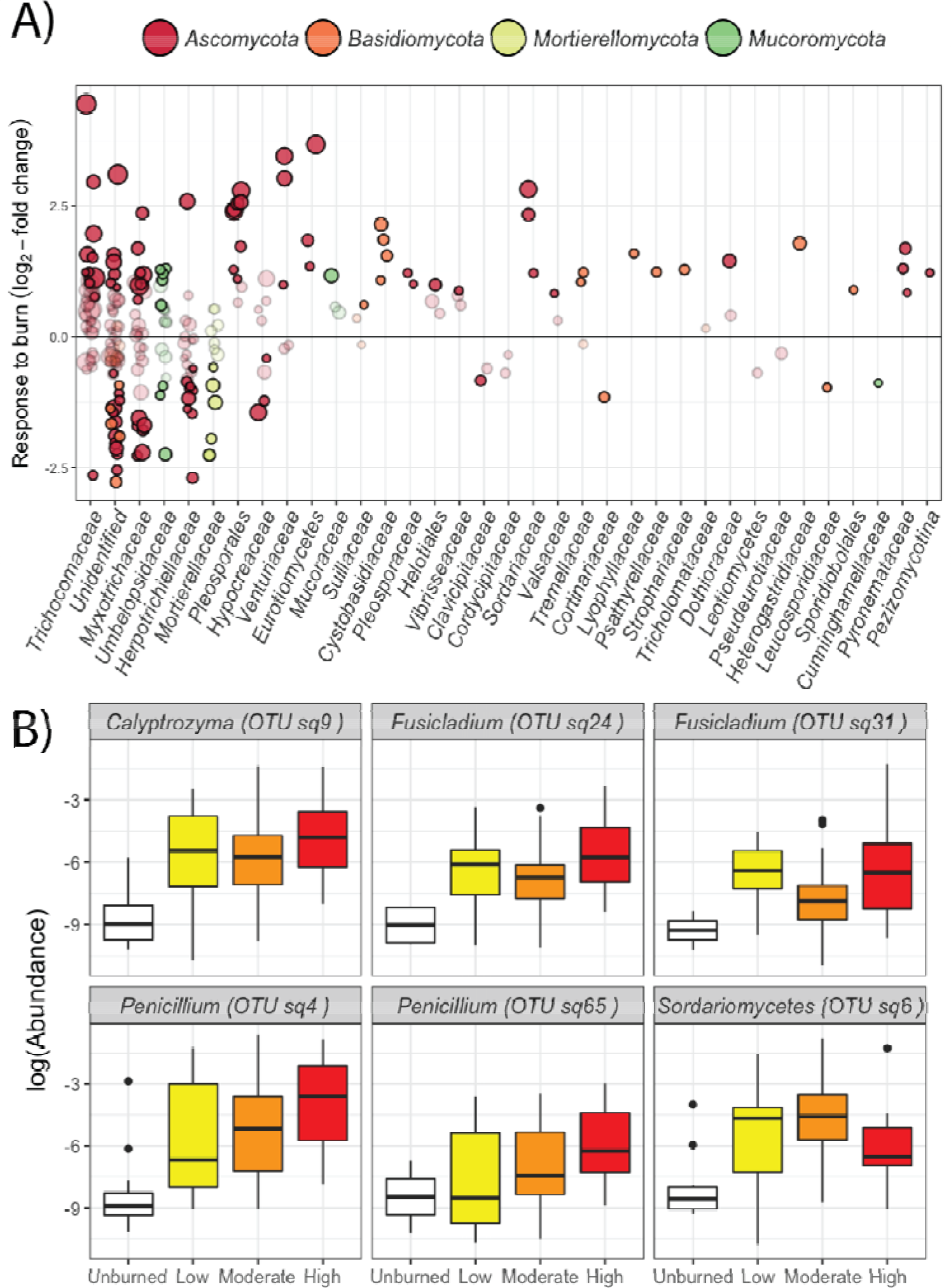
Fungal response to fire. (A) Log_2_-fold change in burned *vs*. unburned plots, controlling for vegetation community, total C, and pH, arranged by class and coloured by phylum. Each point represents a single ITS2 OTU, and the size of each point represents the mean relative abundance of that OTU across all samples. Faint points represent OTUs that were not significantly different in abundance in burned *vs*. unburned plots. (B) Relative abundance (note log scale) of selected fire-responsive fungal OTUs across BSI ranges (unburned=0, 0-2 low, 2-3 moderate, 3-4 high).

### Co-occurrence network

The bacterial and fungal consensus network included 3454 edges or connections, 351 bacterial OTU nodes (~2% of all bacterial OTUs), and 250 fungal OTU nodes (~4% of all fungal OTUs) (Figure 6A, Supplemental Figures 21-26, Supplemental Tables 8 and 9). However, the OTUs that were retained represented an average of 40% (maximum of 74%) of the community for bacteria and 30% for fungi (maximum of 70%). Of the bacteria in the network, 48% were at least a 97% ID match for one of the globally abundant phylotypes designated by Delgado-Baquerizo *et al*. [45] (Supplemental Figure 26). The network had a modularity of 0.58, which is above the threshold of 0.4 suggested to define modular structure [69]. We report the properties of the 8 largest modules in Supplemental Table 9. Two of the larger modules (1 and 11) contain numerous OTUs that were identified as fire-responders using the log_2_-fold-change approach (55% and 82% of OTUs in the module, respectively), while one of the other larger modules (4) is characterized by negative fire-responders (63%). Module 4 also contains OTUs that were prevalent in wetter sites (Figure 6C). Module 11 contains OTUs more prevalent at high pH sites, while Module 1 is characterized OTUs that are more prevalent at low pH sites (Figure 6B). The networks constructed for O horizons or mineral horizons including plants are described in Supplemental Tables 8 and 9. They exhibited similar broad clustering patterns (Supplemental Figures 25-34). Only 1-3 understory plant nodes (*Salix*, *Carex*, and *Geranium*) were retained in the plant consensus networks.

**Figure 6.**
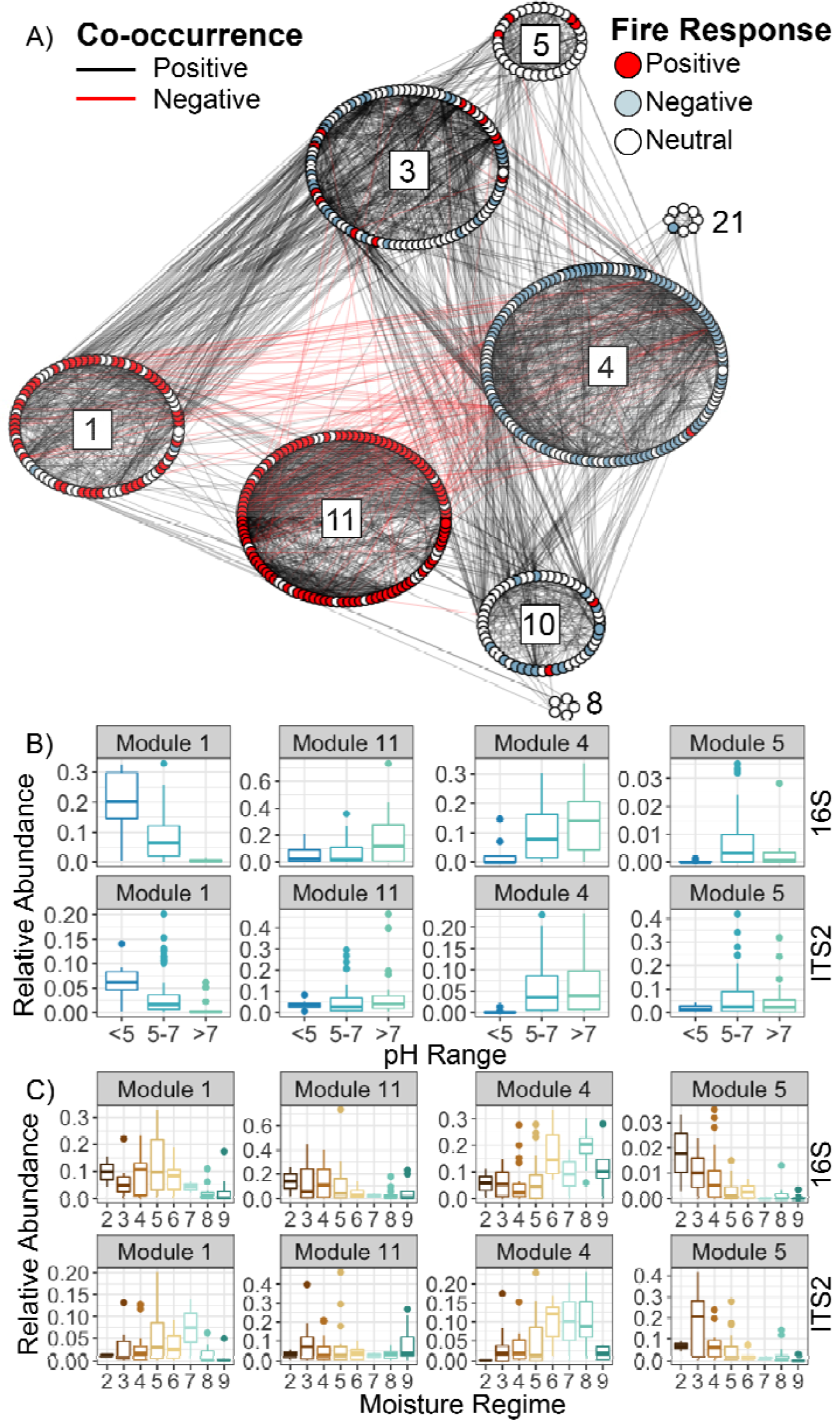
A) Co-occurrence network [16S and ITS2 - Organic and Mineral Horizons], arranged into greedy clustering-defined modules. Each point represents an OTU. Points are coloured by whether they were identified as being significantly more abundant in burned samples (red) and those significantly less abundant in burned samples (light blue) or no significant response (white). Lines between points indicate co-occurrences (black) or co-exclusion (red). Module IDs are indicated with numbers for reference. B) Module representation across moisture regimes: Fraction of total community represented by all bacterial (top, 16S) and fungal (bottom, ITS2) OTUs within selected modules, grouped by moisture regime, 2 being very dry, and 9 being very wet. C) Module representation across pH values: Fraction of total community represented by all bacterial (top, 16S) and fungal (bottom, ITS2) OTUs within selected modules grouped by pH range.

## Discussion

### Burn severity metrics are significant predictors of microbial community composition

Given the wide range of soil properties and vegetation communities spanned by our study, we were impressed that the effects of burning on soil microbial communities stood out so clearly (Figure 2). In addition to the effects of fire, our observation that soil bacterial communities are more strongly structured by pH (than C), while C is a stronger predictor (than pH) for soil fungal communities, is consistent with previous findings [68,70] (Supplemental Note 2). High-throughput sequencing data from this region are rare, but Masse *et al.* [71] and Turney *et al.* [72] report broadly similar microbial communities in northern Alberta to those observed in this study (Supplemental Figures 1-4). However, the novelty of this study lies not only in characterizing the soil microbial communities of - and the effects of fire in - the northern boreal forest in Canada across a very wide range of conditions. The inclusion of burn severity is an important but often “missing piece” in assessing the ecological effects of wildfires, which can range from barely-detectable light surface burns to total tree mortality and complete O horizon losses.

Our data suggest that the predicted increases in burn severity for the boreal forest [18–19] may be accompanied by increasingly disturbed microbial communities (Figure 3A and 3B), although there is little differentiation between different burn severity metrics for predicting microbial community composition. We observed different response trends with severity for different taxa (Figures 4B and 5B), suggesting that fires evoke non-linear responses with increasing severity, and that the shapes of these responses differ for different fire-responsive taxa. Future studies could be designed explicitly to investigate these nonlinear relationships between community composition and burn severity, possibly developing response-based severity categories. Additionally, the study’s sampling timeline (one year post-fire) will affect our detection of different community responses to different severity levels [73]. For example, if the strongest effects of fires on soil microbes are driven by changes to the vegetation community [16], these effects may continue to emerge many years post-fire, while the strongest short-term effects of direct killing of microbes from the fire’s heat may no longer be detectable one-year post-fire. Furthermore, burn severity metrics – originally developed for plant communities – integrate effects that are not as relevant to microbes as to plants, diluting their efficacy as predictors of microbial community composition. Future investigations could decompose the sub-components of burn severity (*e.g.*, degree of understory vegetation survival *vs*. magnitude of combustion of the organic horizon), to determine which are most influential on microbial community composition, and perhaps develop microbially-specific burn severity metrics.

### Specific bacterial and fungal taxa can be identified as fire-responders

Globally abundant organisms that we identified as positive fire responders are likely relevant across diverse ecosystems. For example, Fernández-González *et al*. [74] also observed that both *Arthrobacter sp.* and *Blastococcus sp.* were significantly enriched in post-fire soils, but in a very different ecosystem – oak forests in the Sierra Nevada of Spain. Of their 55 sequenced strains of *Arthrobacter*, 41 isolates were a 100% ID match for our first *Arthrobacter* (OTU sq1) and 11 were a 99% ID match for the second (OTU sq7) [74]. Fernández-González *et al*. [74] speculate that *Arthrobacter* may be able to survive fires due to its ability to resist starvation, desiccation and oxidative stress [25, 75–77]. Then, it may thrive on the fire-affected aromatic C sources [78] and may also play a role in post-fire nitrogen cycling [79] and phosphorus solubilization. These activities could have important effects on plant growth: Fernández-González *et al*. [74] demonstrated 40% or greater increases in plant biomass in alfalfa plants inoculated with a subset of their *Arthrobacter* strains. Thus, our results support their suggestion that *Arthrobacter* may play an important role in the post-fire microbial ecosystem and expand their findings to a very different ecosystem – the boreal forest.

Our most abundant fungal fire-responder likely also has broad ecological relevance. Ten *Penicillium sp.* OTUs were identified as positive fire-responders (Supplemental Table 7), including two that were particularly abundant (OTUs sq4 and sq65; Figure 5B). *Penicillium* is a common saprotrophic forest microfungus [80], and may be taking advantage of the post-fire nutrient and C availability. Mikita-Barbato *et al*. [81] also noted a *Penicillium sp.* that was found at severely burned pine-oak forests in New Jersey, USA, but was not detected at the unburned sites. The increased abundance of *Penicillium* at burned sites could have important ecological consequences: in a global meta-analysis, Bahram *et al*. [68] suggested that increasing fungal antibiotics were associated with an increase in antibiotic resistance genes (ARGs), particularly in association with *Penicillium sp.* Thus, if fires increase *Penicillium* abundances, we might ask, do total bacterial abundances decline and/or do we see an increase in bacteria that may carry ARGs? In our dataset, there is not a significant relationship between the total abundance of *Penicillium* and copies of bacterial 16S genes or bacterial 16S : fungal 18S abundance ratios. However, there are significant positive relationships between the relative abundances of *Penicillium* and the bacterial phylum *Actinobacteria* (p=0.001) and the genus *Streptomyces* (p<0.001). Still, just as the Bahram *et al*. [68] study is correlative, so is this study – it is just as possible that the same underlying factors are increasing both *Penicillium* as well as *Actinobacteria* or *Streptomyces*, rather than that the fire-induced increase in *Penicillium* is somehow selecting for those bacterial taxa.

In addition to the two taxa discussed above, many of the fire-responsive genera we identified have previously been identified as being enriched post-fire in other studies of fungi (*e.g.*, *Neurospora sp.* [82] or *Geopyxis sp.* [83]; Supplemental Note 4) or bacteria [25,79,81,84–85] (Supplemental Note 5). Just like well-established fire-response strategies for plants, there are likely a series of fire-response strategies for microbes (conceptual model illustrated in Figure 7). The first possible trait – fast growth post-fire (as suggested by significantly higher mean predicted 16S gene copy numbers for communities from more severely burned sites, and within positive fire-responders) – may allow a microbe to take advantage of a habitat newly depleted of competitors. Some of the strongest fire-responders (Figure 4) have particularly high predicted copy numbers – e.g., *Massilia sp.* (OTU sq2) has a predicted copy number of 7, while the *Arthrobacter sp.* (OTU sq1) has one of 5.71. This trait has been previously associated with the bacteria that make up early-successional communities, including another post-fire system [58]. It has been suggested that this trait may allow bacteria to grow more quickly [86–87], allowing them to rapidly take advantage of post-fire resources.

**Figure 7.**
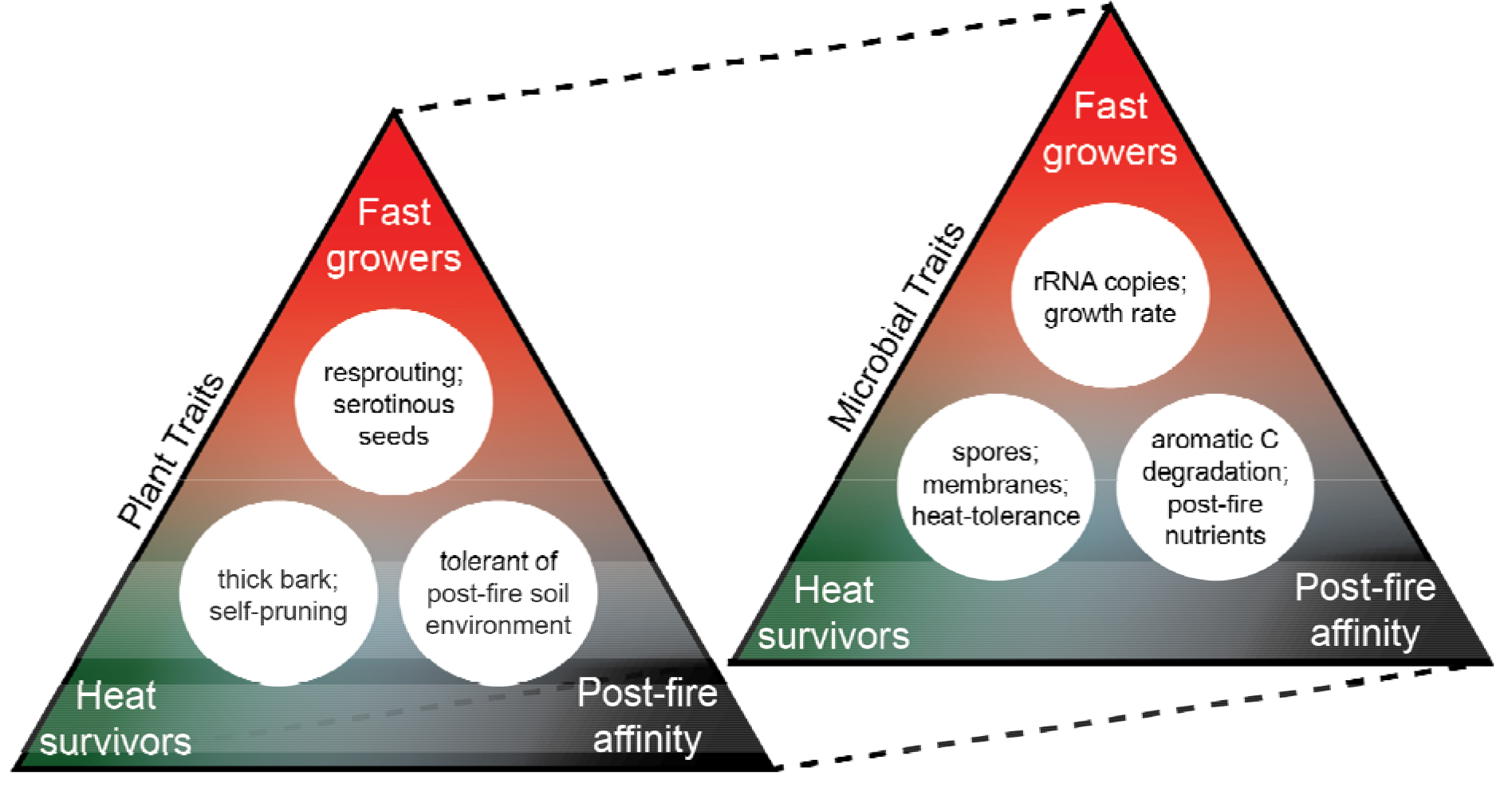
Conceptual figure of hypothesized parallels between fire response strategies for plants and microbes. Layered on top of these traits would be ecological interactions between organisms in the post-fire community.

A second trait that could allow microbes to thrive post-fire would be the ability to exploit resources created by the fire – for example, changes in nutrient availability [88] (Supplemental Note 6) or fire-affected organic matter, which is characterized by an increased abundance of aromatic C structures [89]. Many of our most abundant fire-responsive bacterial taxa (*Aeromicrobium*, *Massilia*, and *Burkholderia-Paraburkholderia*) are genetically identical in the sequenced region to organisms that have been identified as aromatic C-degraders [85,90–91] (Supplemental Note 7). Similarly, numerous fungi with lignolytic capabilities are also able to degrade other aromatic C structures, and include species within the genera *Penicillium* and *Mucor*, for which we identified positive fire-responsive OTUs [34,92] (Figure 5; Supplemental Table 7).

A third potential fire responder trait is survival at elevated temperatures [93–94] (although increased temperatures from fire rapidly attenuate with soil depth [95]). In a study of tree bark-associated fungi in Australia, three *Penicillium spp.* were isolated that could withstand temperatures above 105 °C for one hour, and the authors also noted that teleomorphic taxa (taxa in the sexual reproductive stage) were only found in the heat-treated bark, suggesting that the heat of fires may revive ascospores from dormancy [96]. Because we sampled sites one-year post-fire, the longer-term effects of fire (changes to the soil environment or vegetation) may be playing a larger role than the immediate post-fire effects of the direct killing of organisms during the fire, which might be most important in the weeks or months right after the fire.

In addition to heat survival, fast growth, and the ability to take advantage of post-fire resources, interactions between fire-responders and other members of the ecosystem, including plants and animals, will structure post-fire communities. For example, one OTU identified as a 100% match with *Fimetariella rabenhorstii* was significantly enriched with fire, and has commonly been found in the dung of boreal herbivores [97]. This is consistent with animal studies that suggest that herbivory may increase post-fire as plants regenerate [98–100], and the observation that our study region is inhabited by wood bison that likely benefit from fire clearing grazing areas. With respect to plants, two of the fungal taxa we identified as significant fire-responders were classified as *Fusicladium sp.*, and two as *Phoma sp.* These genera are rated as “probable” and “highly probable” plant pathogens within the FUNGuild classification system [48,101], and one interpretation could be that they are exploiting damaged trees post-fire. However, *Fusicladium* has also been isolated from pine litter [102], and may just as likely be living as a saprotroph on fire-killed litter [103]. Similarly, *Phoma* is also known to exhibit saprotrophic strategies [104]. Other putative plant-associated fungi (pathogenic or non-pathogenic endophytes) were enriched in burned sites (three *Venturia* or *Fusicladium* OTUs). However, we are not able to point to clear broad trends across plant-associated fungal guilds, including ectomycorrhizae, in this dataset (Supplemental Table 7). Additionally, we stress that even 100% ID matches may not have the same functional potential or activity as the organisms identified in reference databases; further study would be required to demonstrate these suggestions. Still, such inter-kingdom interactions merit further study, as they could have implications for the post-fire community assembly in plants and fungi, and the effects of post-fire microbial communities on the broader ecosystem. For example, anecdotally, we have observed patches of jack pine in this region that grow back as unusually dense stands of slow-growing trees after very severe fires. It would be fascinating to determine whether this “stalled growth” could possibly be related to shifts in the microbial community, such as the loss of necessary symbionts.

### Co-occurrence network clusters by fire effects, pH, and moisture regime

The most interesting observation for the network is that the taxa cluster in modules that are associated with fire effects (Figure 6) – the majority of taxa in modules 1 and 11 were independently identified as being positive fire-responders (Figures 4 and 5; Supplemental Tables 6, 7, and 9), many of which are close matches for globally abundant taxa (Supplemental Figure 26). This could prompt future research asking whether the remaining taxa in these modules also respond positively to fire. We also noticed that the two fire-responsive modules (1 and 11) clustered separately – *i.e.*, despite both containing a large proportion of fire-responsive taxa, there are few co-occurrences between the two modules. Our most likely explanation for this is pH: bacterial OTUs from module 1 tend to be more abundant in lower pH soils from across a wide moisture gradient, while bacterial OTUs from module 11 tend to be more abundant in higher pH soils specific to drier ecosystems (Figure 6C). Thus, we might interpret bacterial OTUs in module 11 as broadly representing the high pH fire-responders, and the bacterial OTUs in module 1 as broadly representing the low pH fire-responders.

Many negative fire-responders are captured by Module 4 (Figure 6), which also includes OTUs associated with neutral-high pH (Supplemental Figure 23). This module has a higher abundance in wetlands than other modules do (Figure 6B; Supplemental Table 9), but this largely reflects that wetlands tended to be less severely burned, not that negative fire-responders are generally adapted to wetlands, *per se*. This raises an interesting question of whether wetlands, which tend to burn less severely, play any notable role in seeding the post-fire recovery and reestablishment of microbial communities within the larger, patchwork, landscape. Overall, the network analysis identifies several clusters of fire- and pH-responsive taxa, which could inform future investigations of whether similar patterns are found in different ecosystems that are also affected by fire and to further disentangle the effects of fire on microbes as mediated by changes to vegetation communities and soil properties.

## Conclusions

Despite high cross-site variability, we identified an effect of fire severity on microbial community composition. Building on the efforts of previous studies, our results identify specific fire-responsive microbial taxa, provide support for possible successful post-fire ecological strategies, and suggest that accounting for burn severity could improve our understanding of their response to fires, with potentially important implications for ecosystem functions. Future studies might investigate the most microbially-relevant sub-components of burn severity metrics, continue to classify and test for specific ecological strategies of fire-responsive microbes, establish the timescale over which these effects persist, and determine how prevalent these specific microbial responses to fire are across different ecosystems.

## Acknowledgements

The Government of the Northwest Territories provided in-kind and financial support for the field campaign that produced these data. We also thank Parks Canada Agency and Jean Morin for in-kind support during fieldwork. We acknowledge and thank Xinli Cai, G. Matt Davies, Kathleen Groenewegen, Derek Hall, Koreen Millard, and Doug Stiff for their indispensable assistance in the field. The U.S. Department of Energy award DE-SC0016365 helped support T.W.

## Supplemental Information

Supplemental information files are available online at XXX. All sequences are deposited in the NCBI SRA under accession numbers XXX. Code for the analyses conducted in this paper are available at GitHub.com/TheaWhitman/WoodBuffalo/Paper_Analyses_Figures.

