## Supplemental Information for "Soil Bacterial and Fungal Response to Wildfires in the Canadian Boreal Forest Across a Burn Severity Gradient"

| **Table of Contents** | **Page** |
| --- | --- |
| **Supplemental Figures** |  |
| 1. Relative abundance of dominant bacterial phyla across upland sites. | 4 |
| 2. Relative abundance of dominant bacterial phyla across wetland sites. | 5 |
| 3. Relative abundance of dominant fungal classes across upland sites. | 6 |
| 4. Relative abundance of dominant fungal classes across wetland sites. | 7 |
| 5. NMDS ordination of Bray-Curtis distances between bacterial/archaeal (16S) communities coloured by vegetation community | 8 |
| 6. NMDS ordination of Bray-Curtis distances between fungal (ITS2) communities coloured by vegetation community | 9 |
| 7A. Mean burn severity index (BSI) of sites at which each fungal OTU occurs  7B. Mean moisture regime rating of sites at which each fungal OTU occurs | 10 |
| 8. Mean burn severity index (BSI) of sites at which each fungal OTU occurs, stratified by moisture regime rating and grouped by phylum | 11 |
| 9. Bray-Curtis dissimilarity to unburned sites *vs*. burn severity index of burned sites | 12 |
| 10. Vegetation community dissimilarity vs. organic horizon bacterial and fungal community dissimilarity in uplands and wetlands for all sample pairs | 13 |
| 11. Relationship of log (bacterial 16S gene copy number : fungal 18S gene copy number) with burn severity index (BSI) | 14 |
| 12. Relationship of log (bacterial 16S gene copy number : fungal 18S gene copy number) with pH | 14 |
| 13. Bacterial and archaeal richness estimates | 15 |
| 14. Fungal richness estimates | 15 |
| 15. Log_2_-fold change of bacteria in burned *vs*. unburned plots with Long-Term Soil Productivity taxa highlighted | 16 |
| 16. Relative abundance of most abundant negative fire-responsive bacterial OTUs across severity classes | 16 |
| 17. Proportion of OTUs that are at least 97% ID similar to globally-abundant phylotypes identified by Delgado-Baquerizo *et al*. (2018) for fire responders and non-responders | 17 |
| 18. Mean proportion of total community represented by OTUs that are at least 97% ID similar to globally-abundant phylotypes identified by Delgado-Baquerizo *et al*. (2018) for fire responders and non-responders | 17 |
| 19. Log_2_-fold change of fungi in burned *vs*. unburned plots with Long-Term Soil Productivity taxa highlighted | 18 |
| 20. Relative abundance of most abundant negative fire-responsive fungal OTUs across severity classes | 18 |
| 21. Co-occurrence network, arranged by random walk modules, coloured by fungi vs. bacteria/archaea [16S and ITS2 - Organic and Mineral Horizons]. | 19 |
| 22. Co-occurrence network, arranged by random walk modules, coloured by phylum [16S and ITS2 - Organic and Mineral Horizons]. | 20 |
| 23. Fraction of 16S OTUs in each module from different bacterial phyla [16S and ITS2 - Organic and Mineral Horizons] | 20 |
| 24. Fraction of ITS2 OTUs in each module from different fungal classes [16S and ITS2 - Organic and Mineral Horizons] | 21 |
| 25. Co-occurrence network [16S, ITS2, and plants - Organic Horizons], arranged by random walk modules and coloured by fire response; Module abundances by pH; Module abundances my moisture regime | 22 |
| 26. Co-occurrence network [16S, ITS2, and plants - Organic Horizons], arranged by random walk modules, coloured by fungi bacteria/archaea, and plants | 23 |
| 27. Co-occurrence network [16S, ITS2, and plants - Organic Horizons], arranged by random walk modules, coloured by phylum | 24 |
| 28. Fraction of 16S OTUs in each module from different bacterial phyla [16S, ITS2, and plants - Organic Horizons] | 24 |
| 29. Fraction of ITS2 OTUs in each module from different fungal classes [16S, ITS2, and plants - Organic Horizons] | 25 |
| 30. Co-occurrence network [16S, ITS2, and plants - Mineral Horizons], arranged by random walk modules and coloured by fire response; Module abundances by pH; Module abundances my moisture regime | 26 |
| 31. Co-occurrence network [16S, ITS2, and plants - Mineral Horizons], arranged by random walk modules, coloured by fungi bacteria/archaea, and plants | 27 |
| 32. Co-occurrence network [16S, ITS2, and plants - Mineral Horizons], arranged by random walk modules, coloured by phylum | 28 |
| 33. Fraction of 16S OTUs in each module from different bacterial phyla [16S, ITS2, and plants - Mineral Horizons] | 28 |
| 34. Fraction of ITS2 OTUs in each module from different fungal classes [16S, ITS2, and plants - Mineral Horizons] | 29 |
| **Supplemental Tables** |  |
| 1. Site characterization summary | 30 |
| 2. Soil properties | 31 |
| 3. Primers used in this study | 32 |
| 4. 16S Illumina PCR primers (available as .csv) | 32 |
| 5. ITS2 Illumina PCR primers (available as .csv) | 32 |
| 6. Fire-responsive 16S OTUs (available as .csv) | 32 |
| 7. Fire-responsive ITS2 OTUs (available as .csv) | 32 |
| 8. Co-occurrence network properties for full network | 33 |
| 9. Properties of co-occurrence network modules | 34 |
| 10. Connector and hub taxa in co-occurrence network [16S and ITS2 - Organic and Mineral Horizons] (available as .csv) | 34 |
| 11. Connector and hub taxa in co-occurrence network [Plants, 16S, and ITS2 - Organic Horizons] (available as .csv) | 34 |
| 12. Connector and hub taxa in co-occurrence network [Plants, 16S, and ITS2 - Mineral Horizons] (available as .csv) | 34 |
| **Supplemental Notes** |  |
| 1. Full methodological details | 35 |
| 2. Significant predictors of community composition | 41 |
| 3. Fires did not significantly affect richness or fungal:bacterial ratios one year post-fire | 42 |
| 4. Discussion of fire-responsive fungal genera and phyla in other studies | 43 |
| 5. Discussion of fire-responsive bacterial genera in other studies | 44 |
| 6. *Calyptrozyma* as putative nutrient-responsive positive fire-responder | 45 |
| 7. Fire-responder similarity to PAH-degrading organisms | 46 |
| 8. Co-occurrence network and ericoid fungi / ericaceous plants | 47 |
| **References** | 48 |

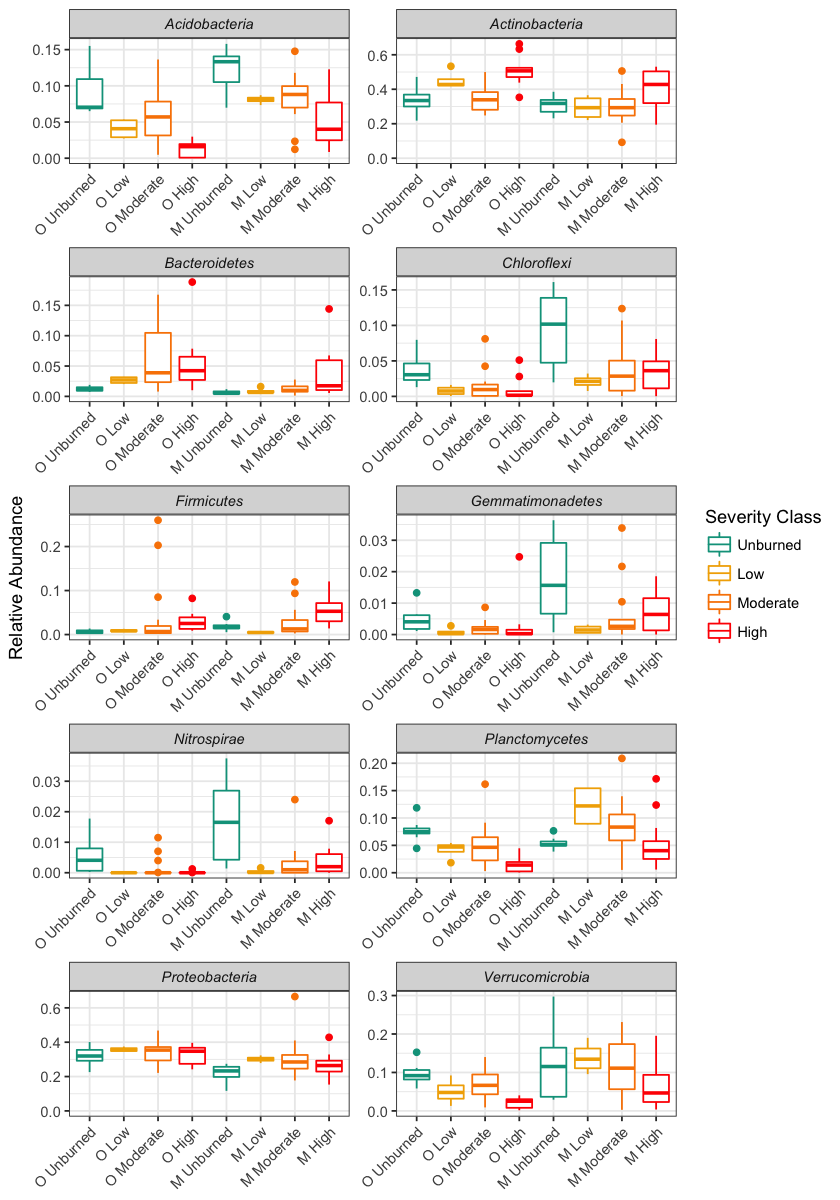

Supplemental Figure 1. Relative abundance of dominant bacterial phyla across upland sites, plotted for organic (“O”) and mineral (“M”) horizons across BSI ranges (unburned=0, 0-2 low, 2-3 moderate, 3-4 high).

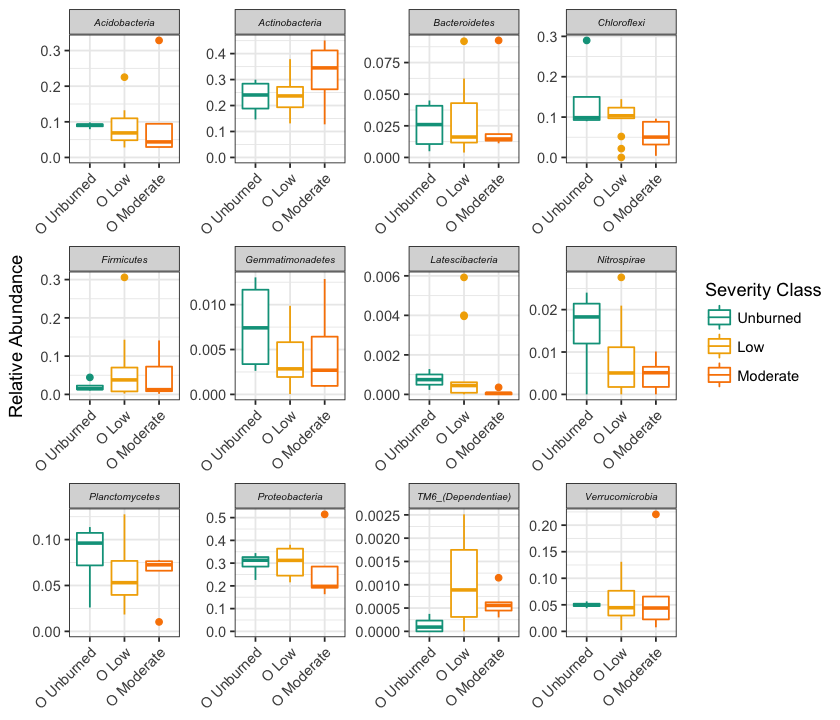

Supplemental Figure 2. Relative abundance of dominant bacterial phyla across wetland sites, plotted for organic (“O”) horizons across BSI ranges (unburned=0, 0-2 low, 2-3 moderate, 3-4 high).

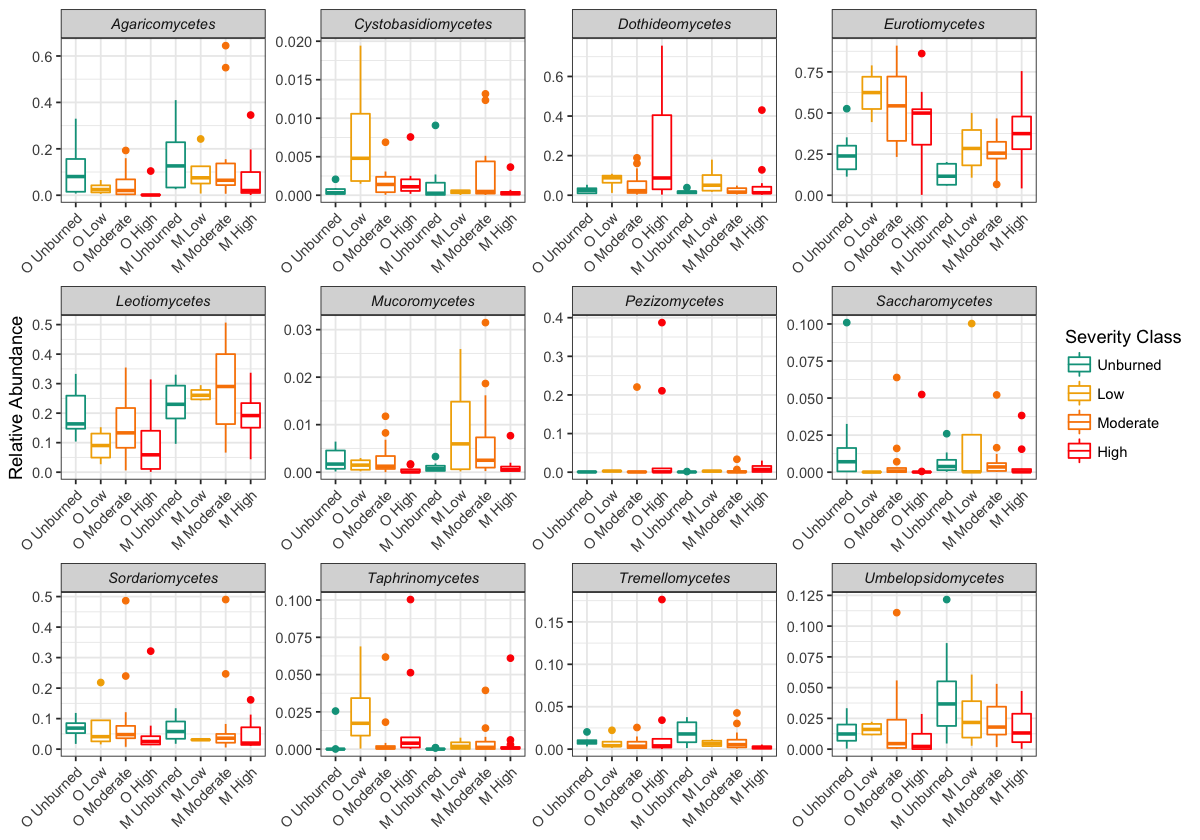

Supplemental Figure 3. Relative abundance of dominant fungal classes across upland sites, plotted for organic (“O”) and mineral (“M”) horizons across BSI ranges (unburned=0, 0-2 low, 2-3 moderate, 3-4 high).

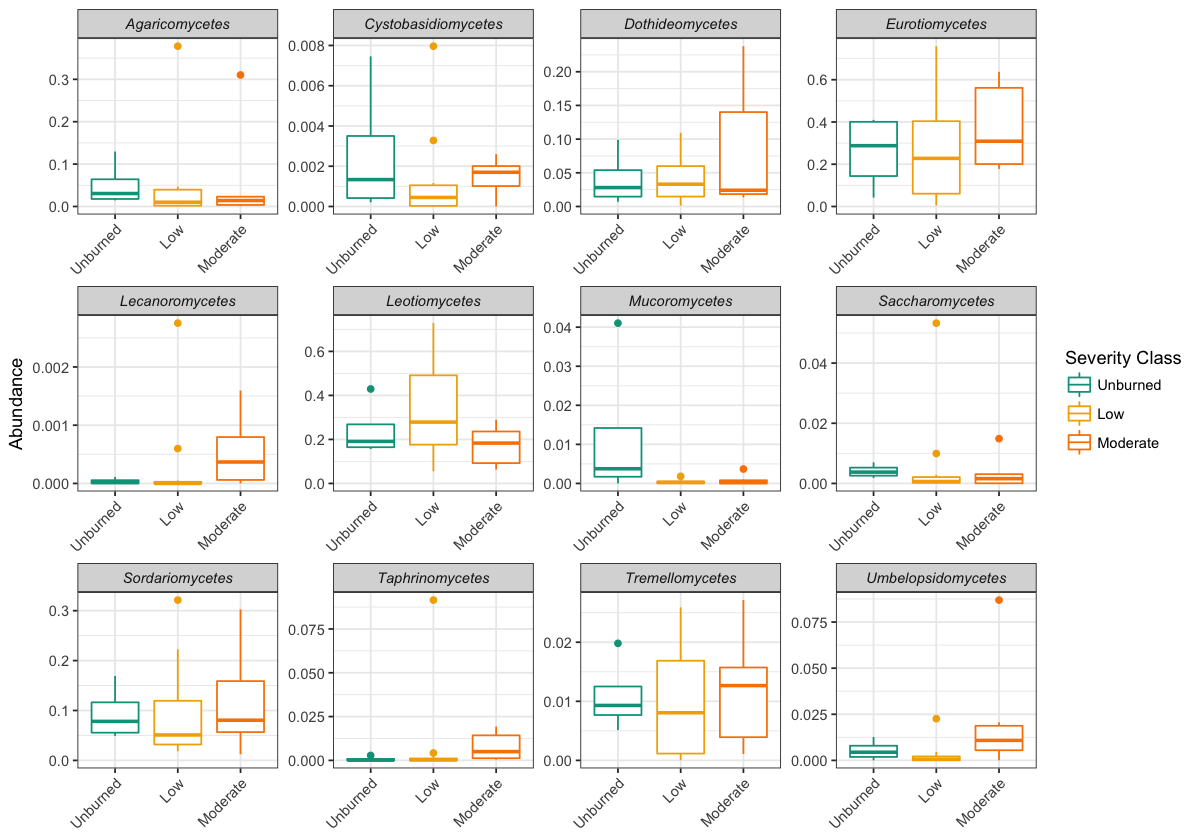

Supplemental Figure 4. Relative abundance of dominant fungal classes across wetland sites, plotted for organic (“O”) horizons across BSI ranges (unburned=0, 0-2 low, 2-3 moderate, 3-4 high).

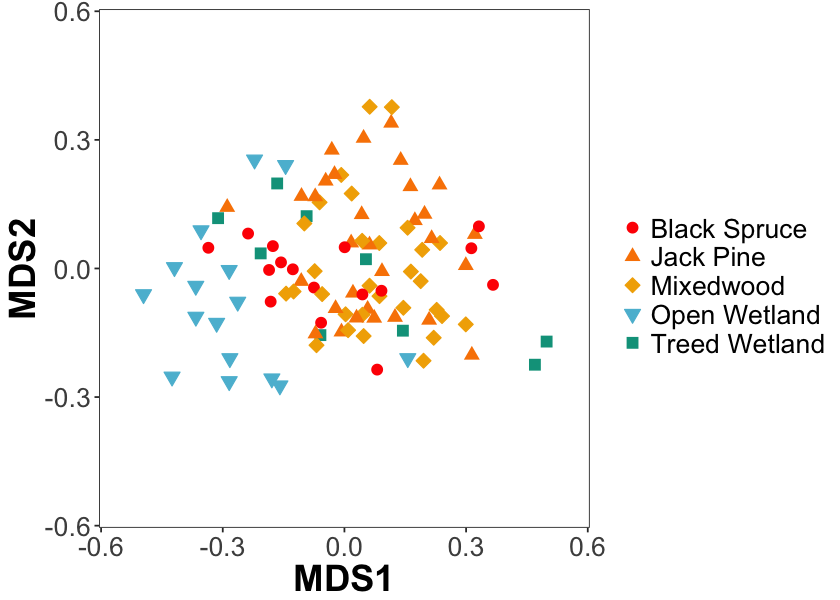

Supplemental Figure 5. NMDS ordination of Bray-Curtis distances between bacterial/archaeal (16S) communities for all samples (k=2, stress=0.16). Points are coded by dominant overstory vegetation community - red circles indicate black spruce, orange upward triangles indicate jack pine, yellow diamonds represent mixedwood, blue downward triangles indicate open wetland, green squares represent treed wetland.

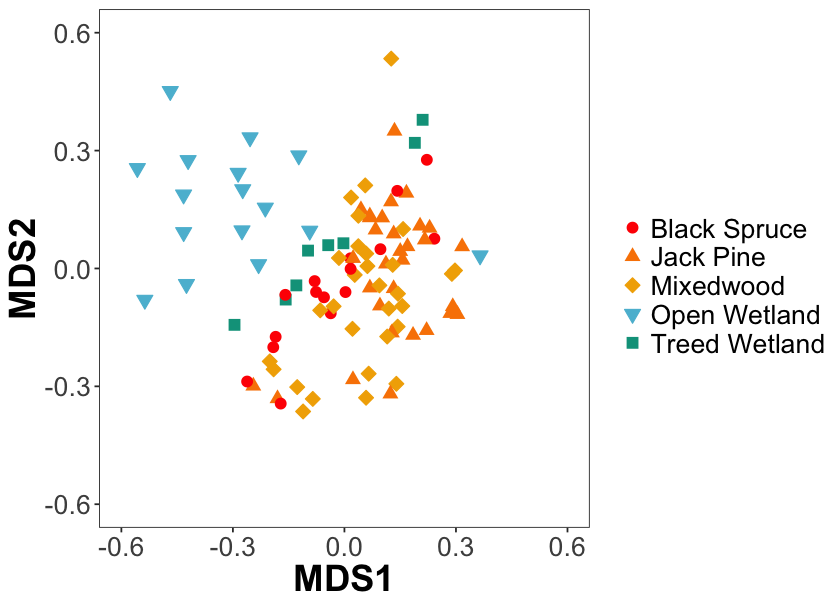

Supplemental Figure 6. NMDS ordination of Bray-Curtis distances between fungal (ITS2) communities for all samples (k=3, stress=0.14). Points are coded by dominant overstory vegetation community - red circles indicate black spruce, orange upward triangles indicate jack pine, yellow diamonds represent mixedwood, blue downward triangles indicate open wetland, green squares represent treed wetland.

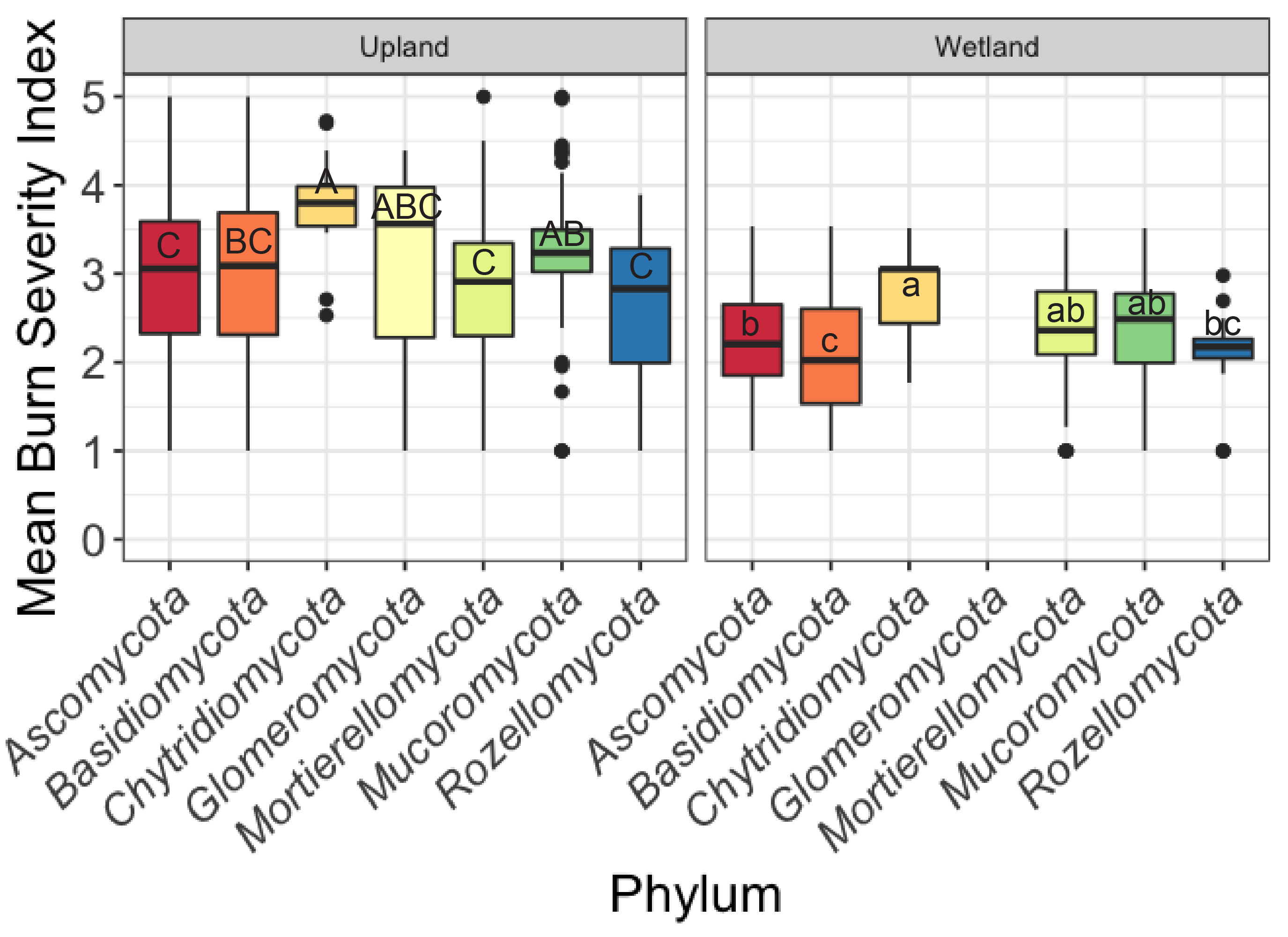

Supplemental Figure 7A. Mean burn severity index (BSI) of sites at which each fungal OTU occurs, grouped by phylum, for upland (left) and wetland (right) sites. Letters indicate significant differences between phyla (ANOVA, Tukey’s HSD, p<0.05).

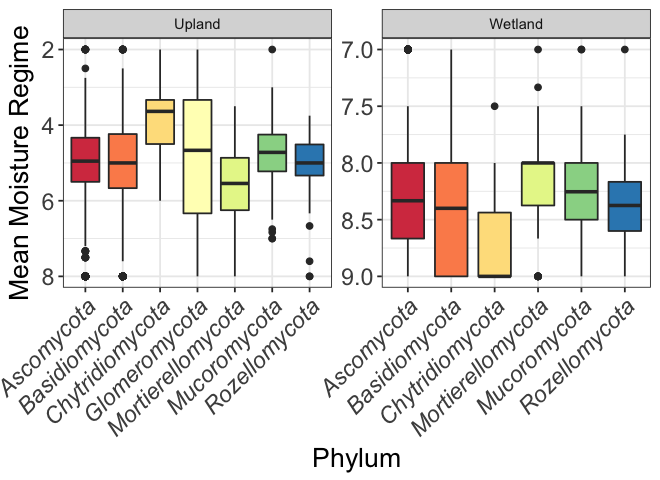

Supplemental Figure 7B. Mean moisture regime rating of sites at which each fungal OTU occurs, grouped by phylum, for upland (left) and wetland (right) sites. (Moisture scale is presented in reverse order to mirror equivalent plot for easier comparison to the companion Composite Burn Index plot in Supplemental Figure 7A.) High values represent wetter conditions, while low values represent drier conditions.

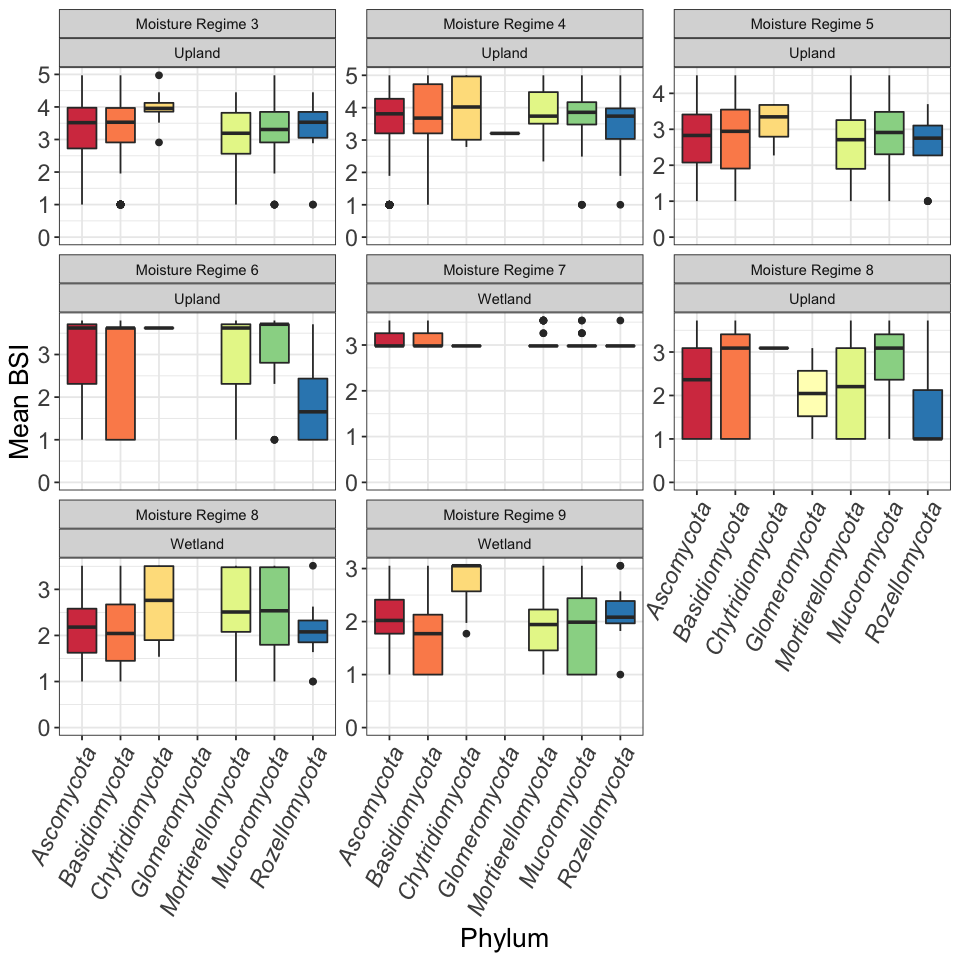

Supplemental Figure 8. Mean burn severity index (BSI) of sites at which each fungal OTU occurs, stratified by moisture regime rating and grouped by phylum.

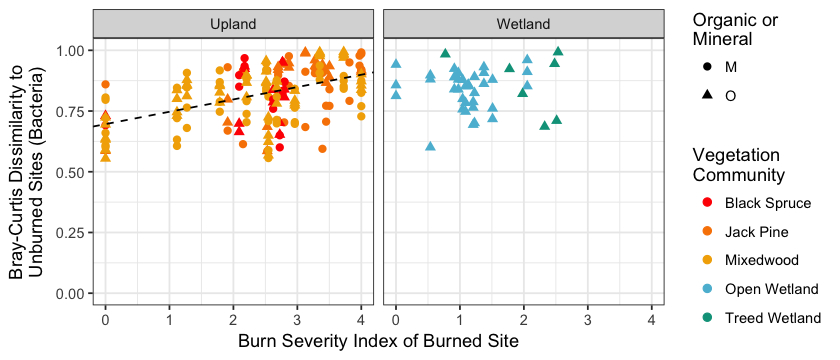

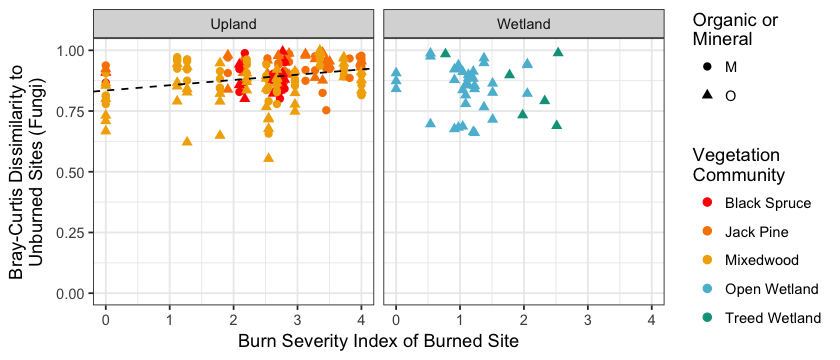

Supplemental Figure 9. Bray-Curtis dissimilarity to unburned sites (within the same vegetation community and the same soil horizon type) for bacteria (top) and fungi (bottom) in uplands (left) and wetlands (right) vs. burn severity index of burned sites. Points are coloured by vegetation community; circles represent mineral horizon samples, triangles represent organic horizon samples. Dashed lines indicate linear regressions (bacteria: y = 0. 0.05 x + 0.70, p<0.001, R^2^_adj_ = 0.22; fungi: y = 0.02 x + 0.83, p<0.001, R^2^_adj_ = 0.08)

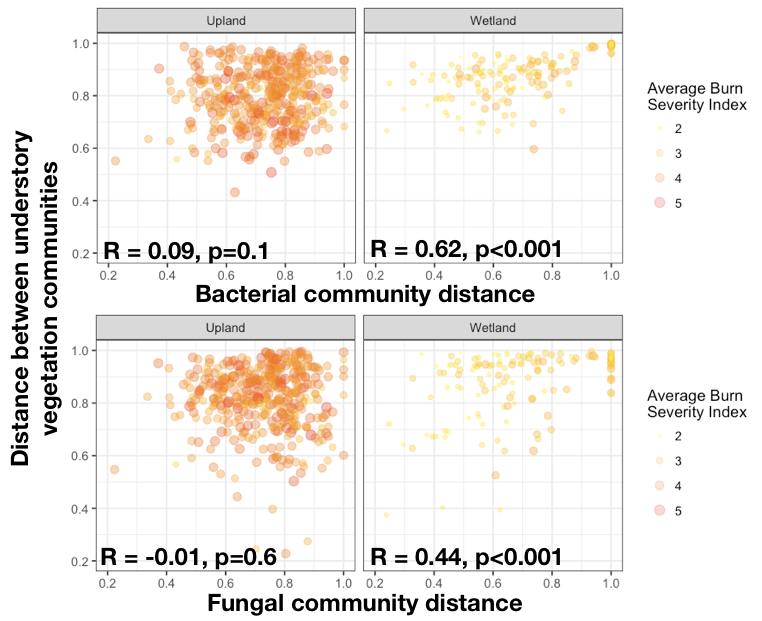

Supplemental Figure 10. Vegetation community dissimilarity vs. organic horizon bacterial (top) and fungal (bottom) community dissimilarity (Bray-Curtis) in uplands (left) and wetlands (right) for all sample pairs. Each point represents a pair of samples; points are coloured by average burn severity index of the pair. There is no significant correlation for upland samples (Mantel tests, 999 permutations). There are significant positive correlations for wetland samples (Mantel tests, 999 permutations).

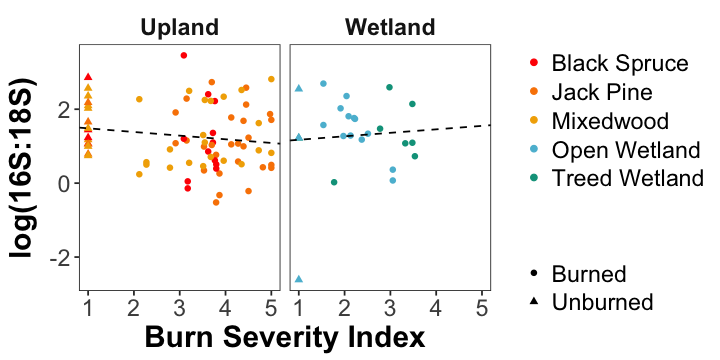

Supplemental Figure 11. Relationship of log (bacterial 16S gene copy number : fungal 18S gene copy number) with burn severity index (BSI). Points are coloured by vegetation community. Triangles indicated burned samples, circles indicate unburned samples. Linear correlations plotted separately for upland sites (p=0.19) and wetland sites (p=0.74).

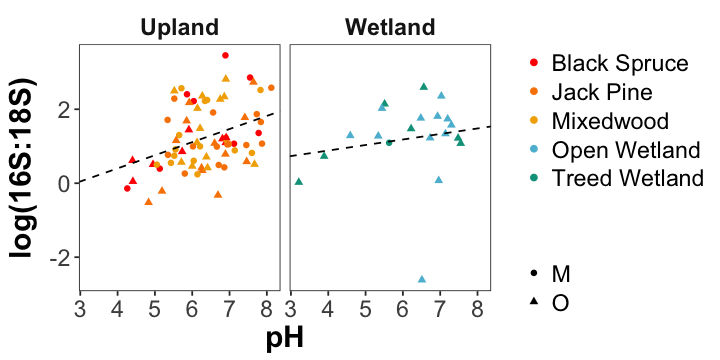

Supplemental Figure 12. Relationship of log (bacterial 16S gene copy number : fungal 18S gene copy number) with pH. Points are coloured by vegetation community. Triangles indicated burned samples, circles indicate unburned samples. Linear correlations for upland sites (R^2^_adj_=0.13, p=0.001) and wetland sites (p=0.49).

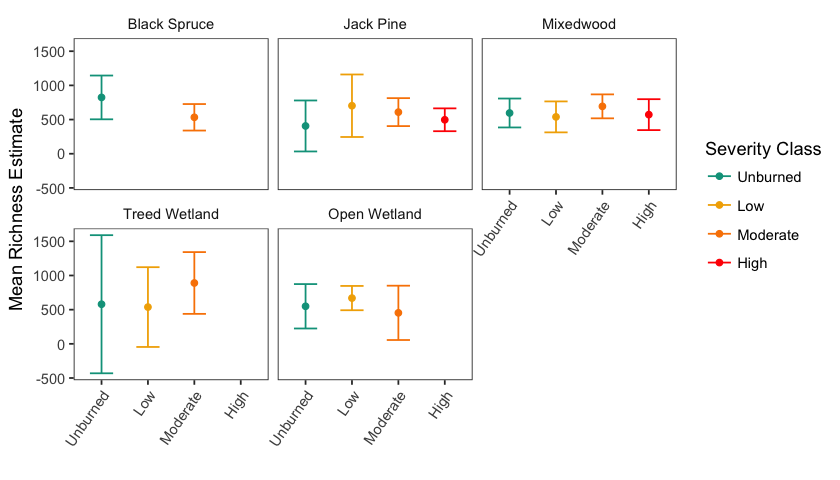

Supplemental Figure 13. Bacterial and Archaeal richness (number of 16S OTUs) estimates for each vegetation community and burn severity index ranges (unburned=0, 0-2 low, 2-3 moderate, 3-4 high), as predicted using best fit model in *breakaway*. Error bars represent ±1.96SE.

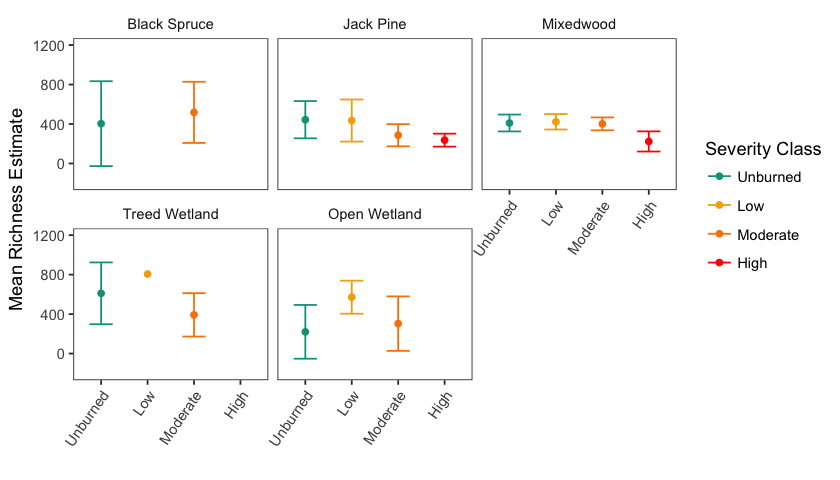

Supplemental Figure 14. Fungal richness (number of ITS2 OTUs) estimates for each vegetation community and burn severity index ranges (unburned=0, 0-2 low, 2-3 moderate, 3-4 high), as predicted using best fit model in *breakaway*. Error bars represent ±1.96SE.

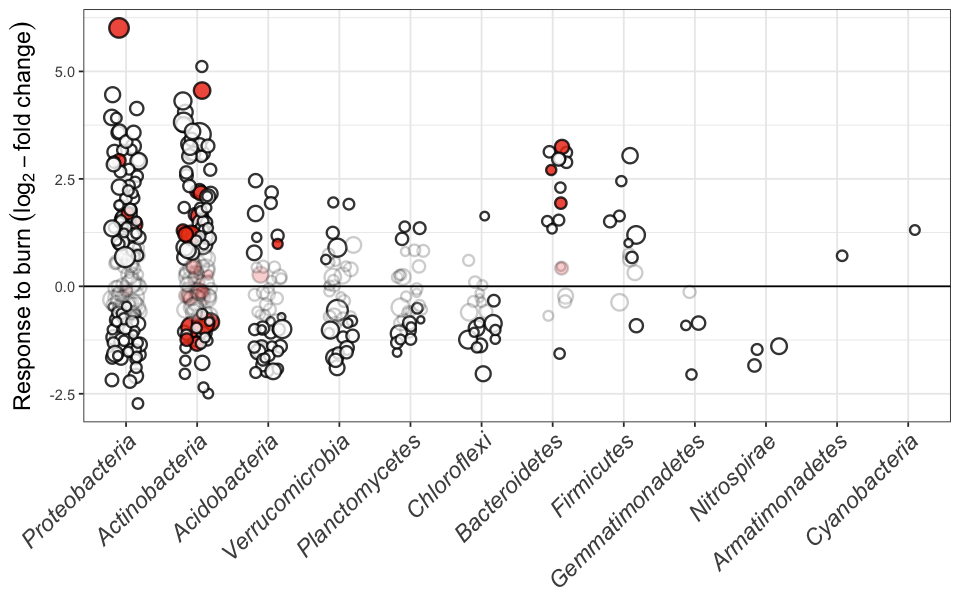

Supplemental Figure 15. Log_2_-fold change in burned *vs*. unburned plots, controlling for vegetation community, total C, and pH. Each point represents a single 16S OTU, and the size of each point represents the mean relative abundance of that OTU across all samples. Red shaded points represent OTUs from genera that were classified as being responsive to organic matter removal by Wilhelm et al. (2017).

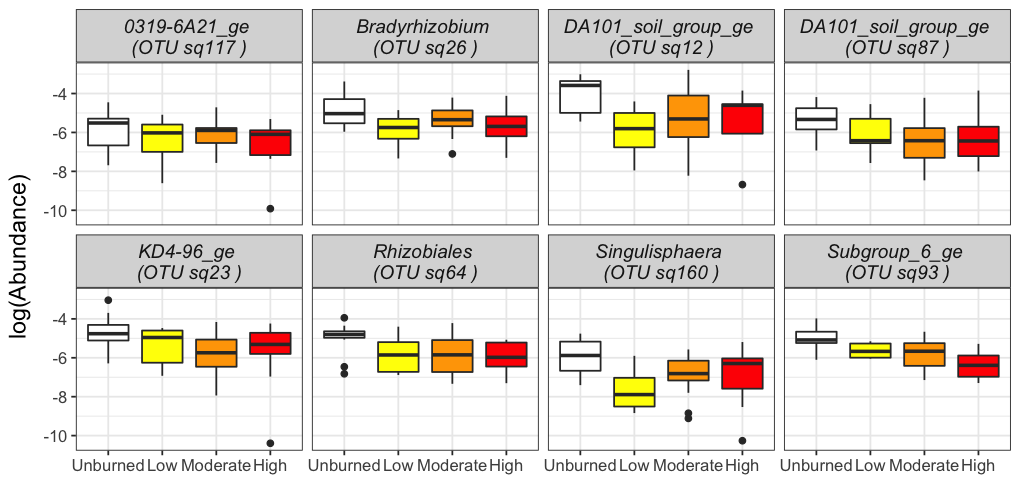

Supplemental Figure 16. Relative abundance (note log scale) of most abundant negative fire-responsive bacterial OTUs across BSI ranges (unburned=0, 0-2 low, 2-3 moderate, 3-4 high). All OTUs are significantly less abundant in burned vs. unburned soils.

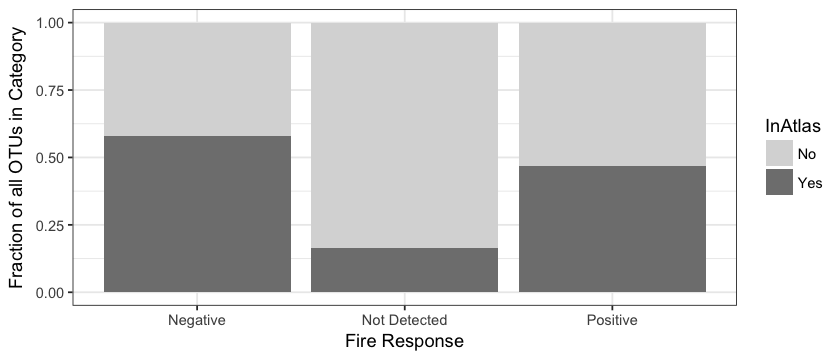

Supplemental Figure 17. Proportion of OTUs that are at least 97% ID similar to globally-abundant phylotypes identified by Delgado-Baquerizo *et al*. (2018) (dark grey bars) for OTUs identified as being negative (n=133) or positive fire responders (n=160), and those not detected as either positively or negatively fire-responsive (n=19,988).

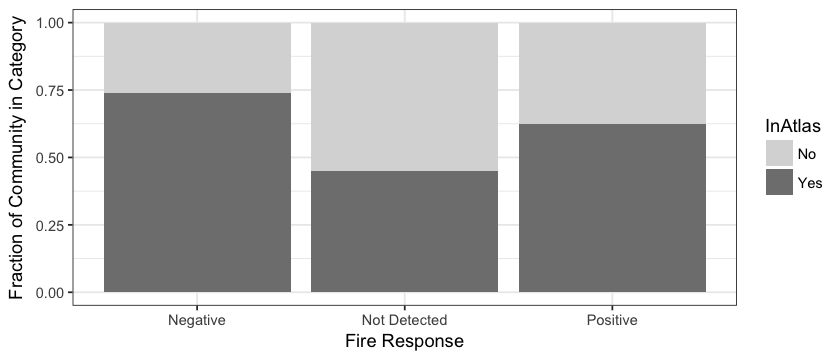

Supplemental Figure 18. Mean proportion of total community represented by OTUs that are at least 97% ID similar to globally-abundant phylotypes identified by Delgado-Baquerizo *et al*. (2018) (dark grey bars) for OTUs identified as being negative (n=133) or positive fire responders (n=160), and those not detected as either positively or negatively fire-responsive (n=19,988), for all samples in this dataset.

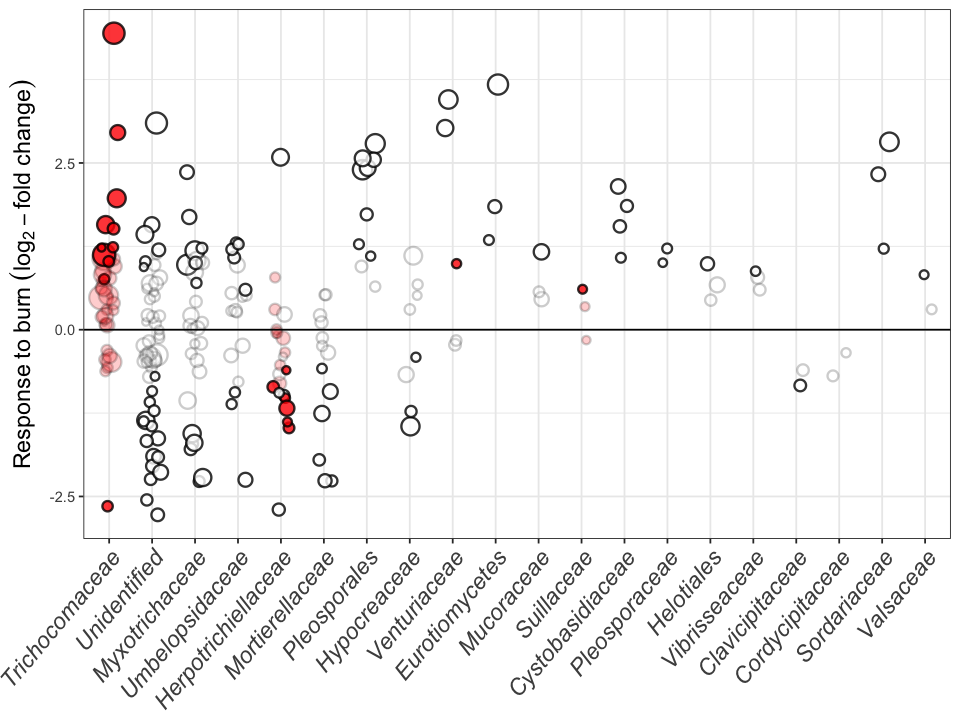

Supplemental Figure 19. Log_2_-fold change in burned *vs*. unburned plots, controlling for vegetation community, total C, and pH. Each point represents a single ITS2 OTU, and the size of each point represents the mean relative abundance of that OTU across all samples. Red shaded points represent OTUs from genera that were classified as being responsive to organic matter removal by Wilhelm et al. (2017).

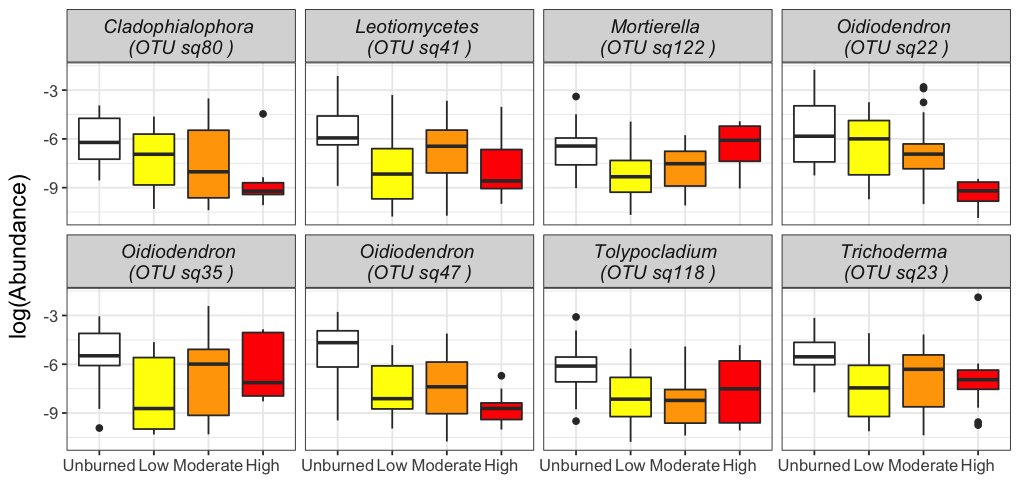

Supplemental Figure 20. Relative abundance (note log scale) of most abundant negative fire-responsive fungal OTUs across BSI ranges (unburned=0, 0-2 low, 2-3 moderate, 3-4 high). All OTUs are significantly less abundant in burned vs. unburned soils. The *Leotiomycetes* OTU is a 100% ID match to a *Gymnostellatospora alpina* (Vu *et al.*, 2019).

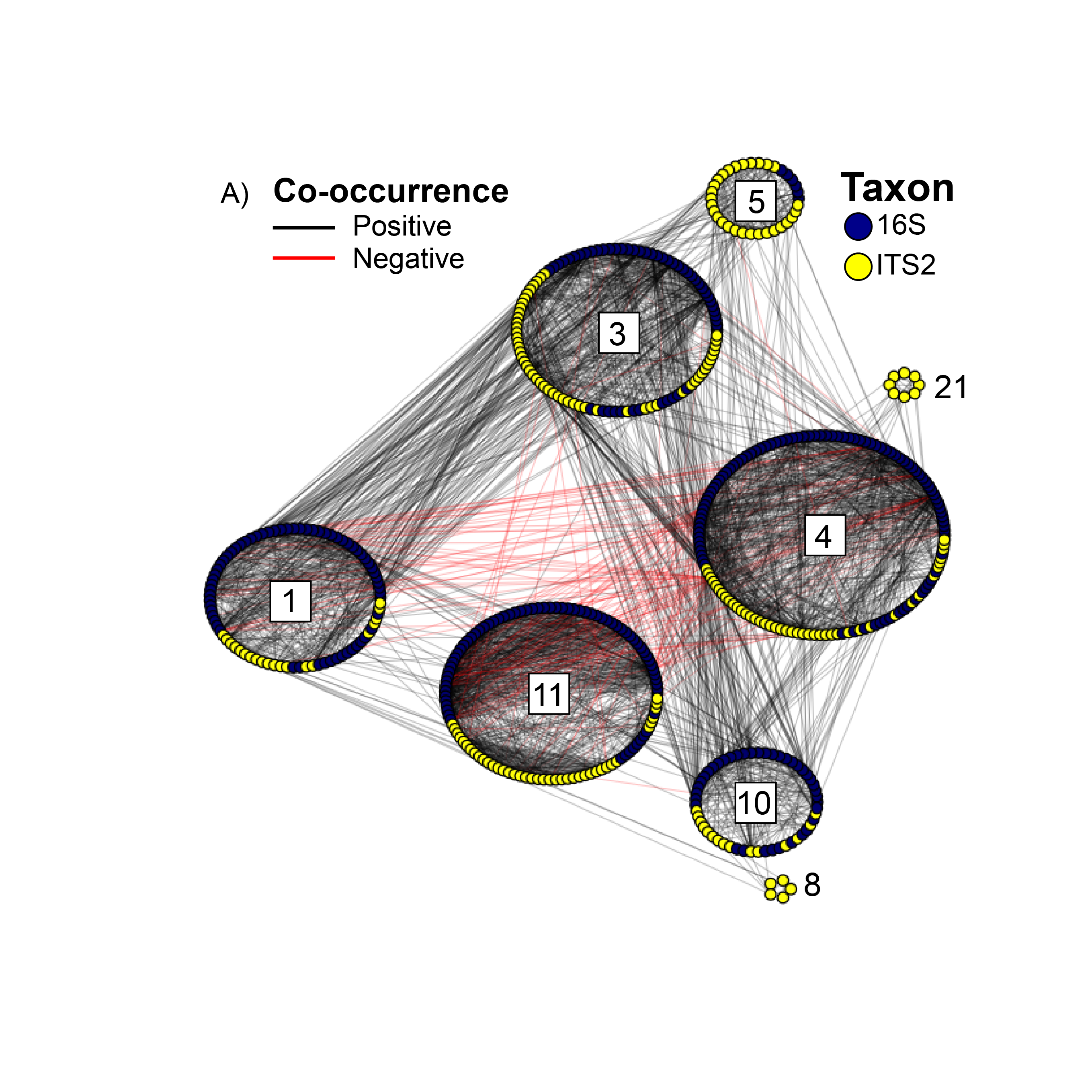

Supplemental Figure 21. Co-occurrence network [16S and ITS2 - Organic and Mineral Horizons], arranged by random walk modules. Each point represents an OTU. Yellow=Fungi (ITS2), Navy=Bacteria and Archaea (16S). Lines between points indicate co-occurrences (black) or co-exclusion (red). Module IDs are indicated with numbers for reference.

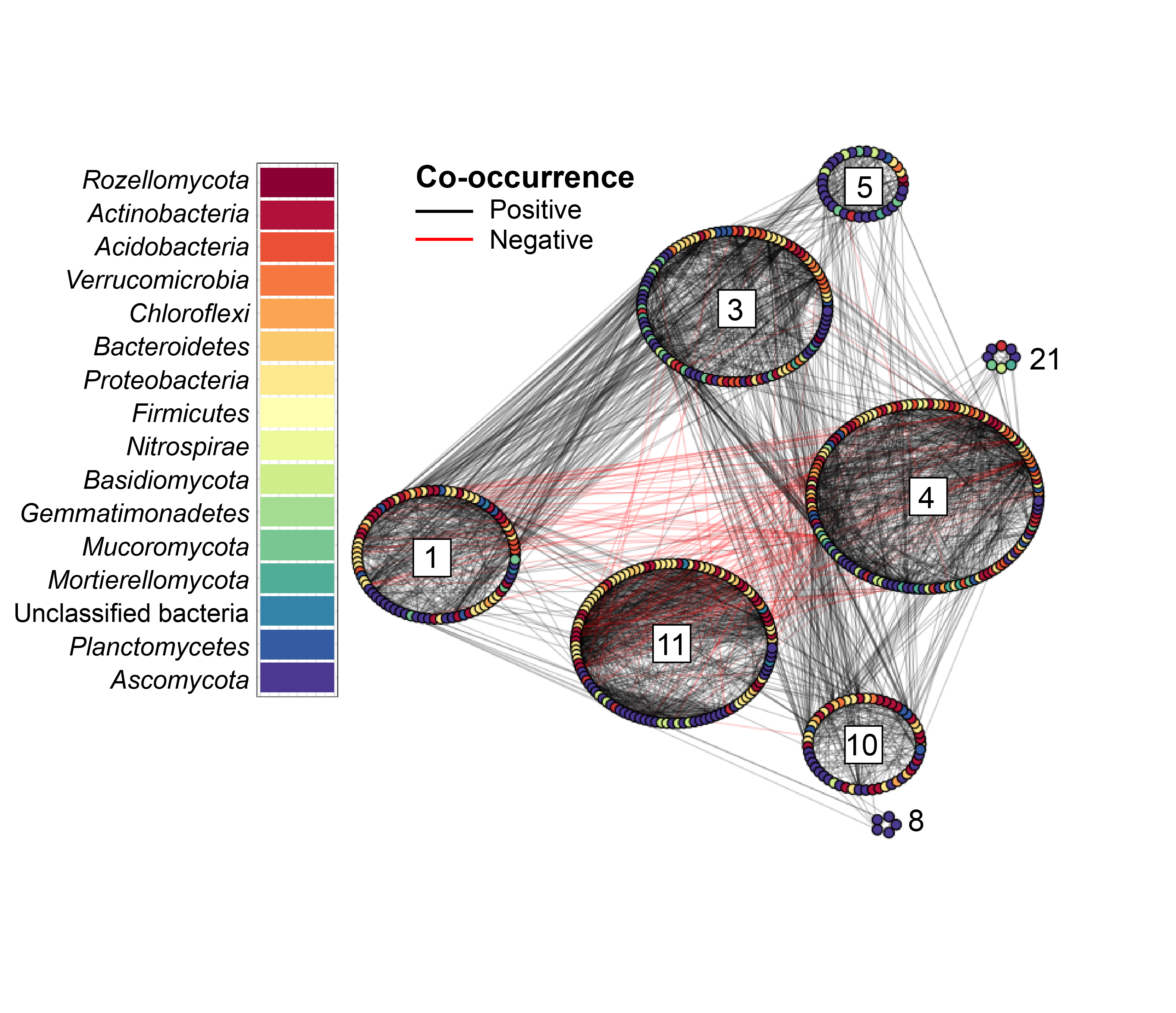

Supplemental Figure 22. Co-occurrence network [16S and ITS2 - Organic and Mineral Horizons], arranged by random walk modules. Each node represents an OTU and is coloured by phylum. Lines between points indicate co-occurrences (black) or co-exclusion (red). Module IDs are indicated with numbers for reference.

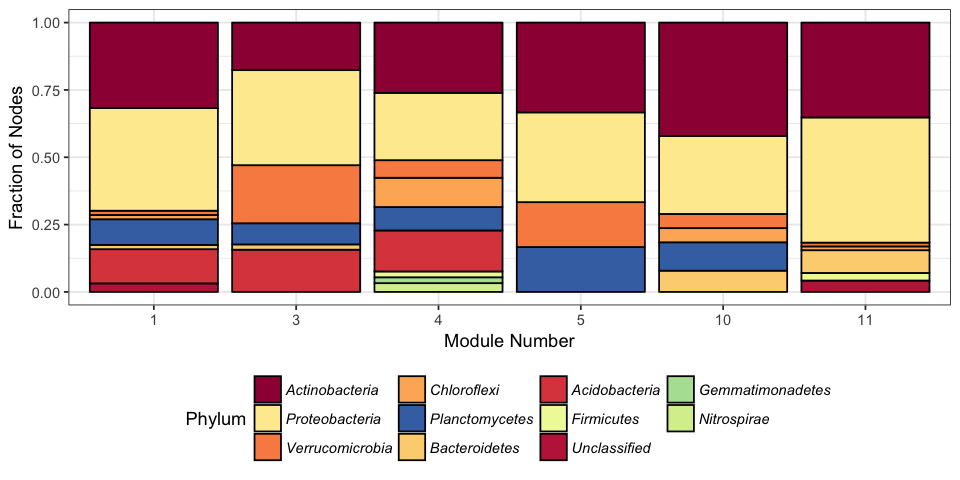

Supplemental Figure 23. Fraction of 16S OTUs in each module from different bacterial phyla in co-occurrence network [16S and ITS2 - Organic and Mineral Horizons].

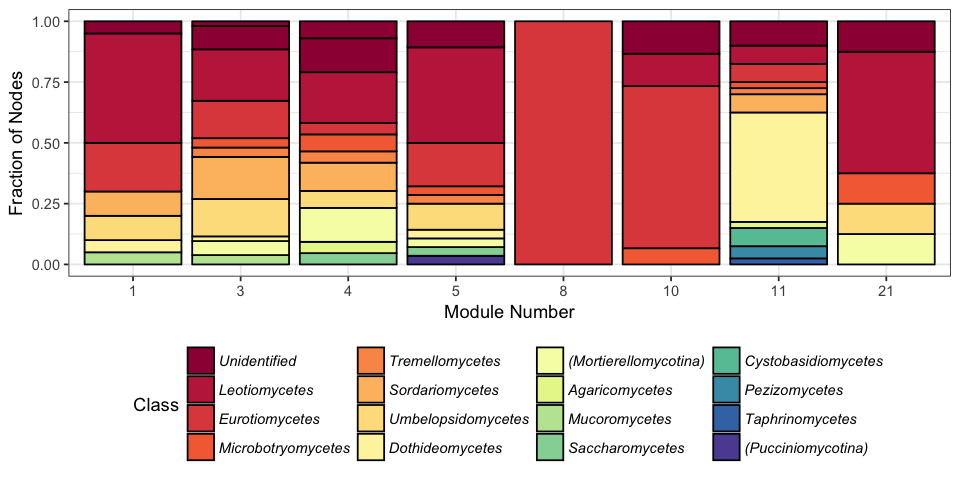

Supplemental Figure 24. Fraction of ITS2 OTUs in each module from different fungal classes in co-occurrence network [16S and ITS2 - Organic and Mineral Horizons].

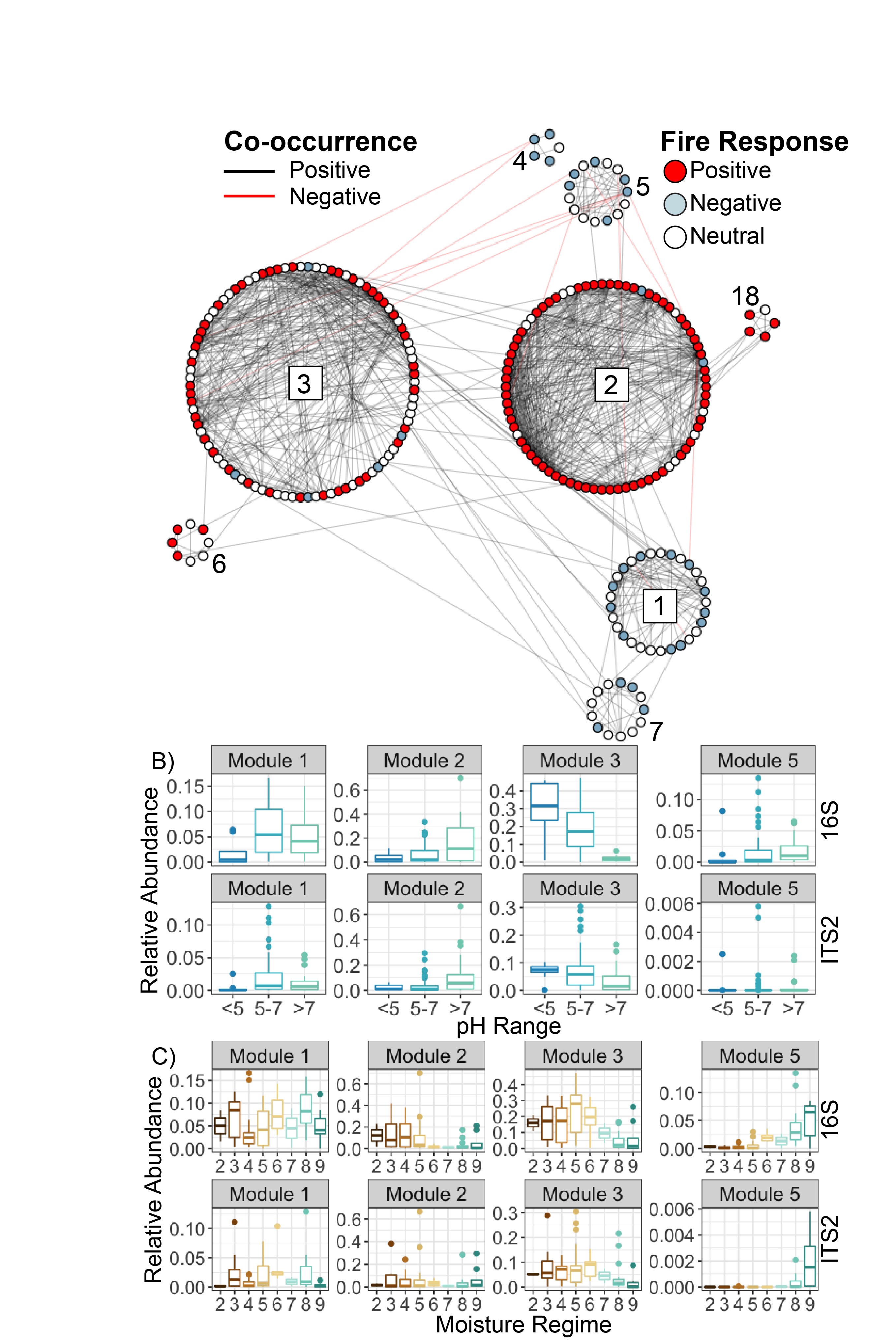

Supplemental Figure 25. A) Co-occurrence network [16S, ITS2, and plants - Organic Horizons], arranged into greedy clustering-defined modules. Each point represents an OTU. Points are coloured by whether they were identified as being significantly more abundant in burned samples (red) and those significantly less abundant in burned samples (light blue) or no significant response (white). Lines between points indicate co-occurrences (black) or co-exclusion (red). Module IDs are indicated with numbers for reference. B) Module representation across moisture regimes: Fraction of total community represented by all bacterial (top, 16S) and fungal (bottom, ITS2) OTUs within selected modules, grouped by moisture regime, 2 being very dry, and 9 being very wet. C) Module representation across pH values: Fraction of total community represented by all bacterial (top, 16S) and fungal (bottom, ITS2) OTUs within selected modules grouped by pH range.

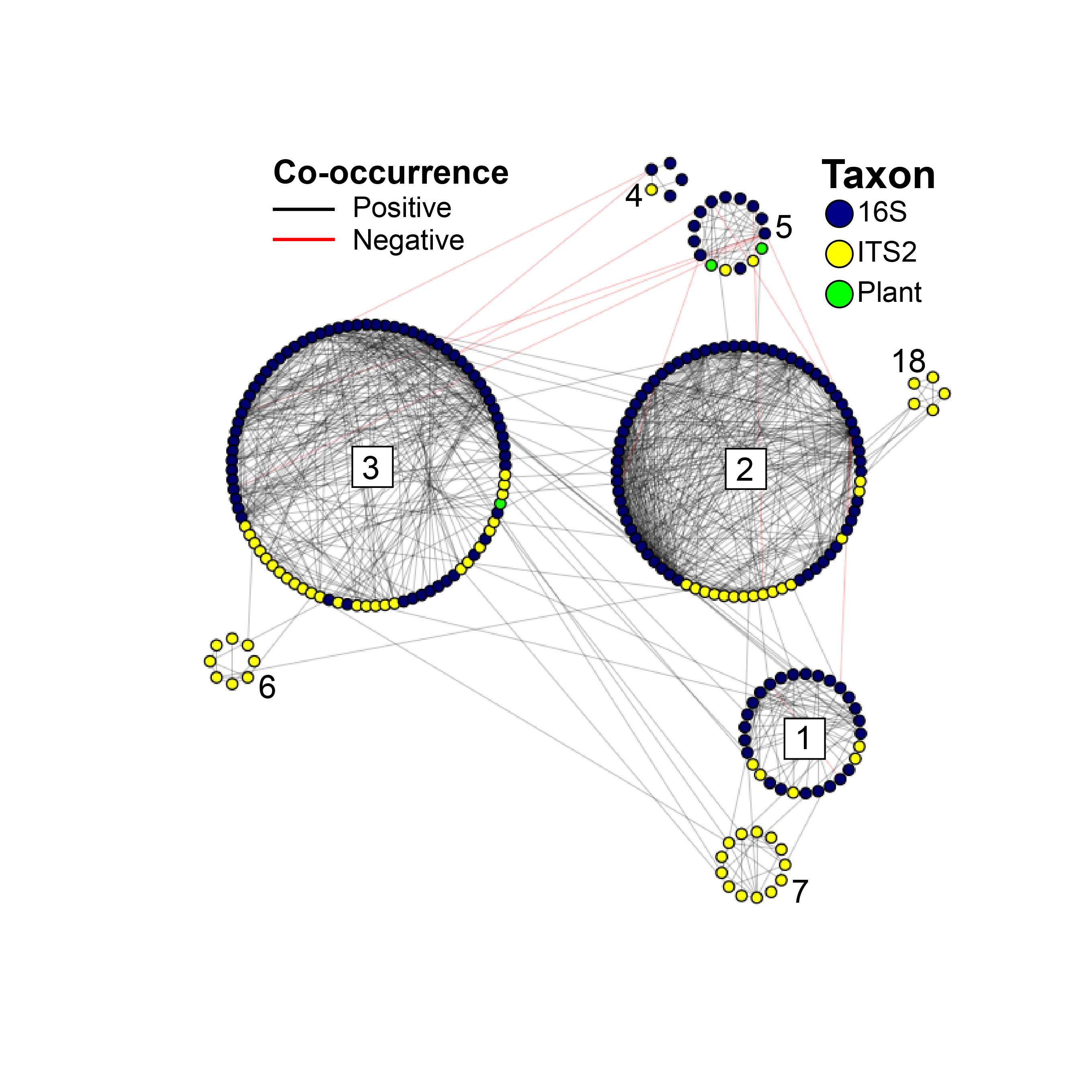

Supplemental Figure 26. Co-occurrence network [16S, ITS2, and plants - Organic Horizons], arranged by random walk modules. Each point represents an OTU. Yellow=Fungi (ITS2), Navy=Bacteria and Archaea (16S), Green= Plants. Lines between points indicate co-occurrences (black) or co-exclusion (red). Module IDs are indicated with numbers for reference.

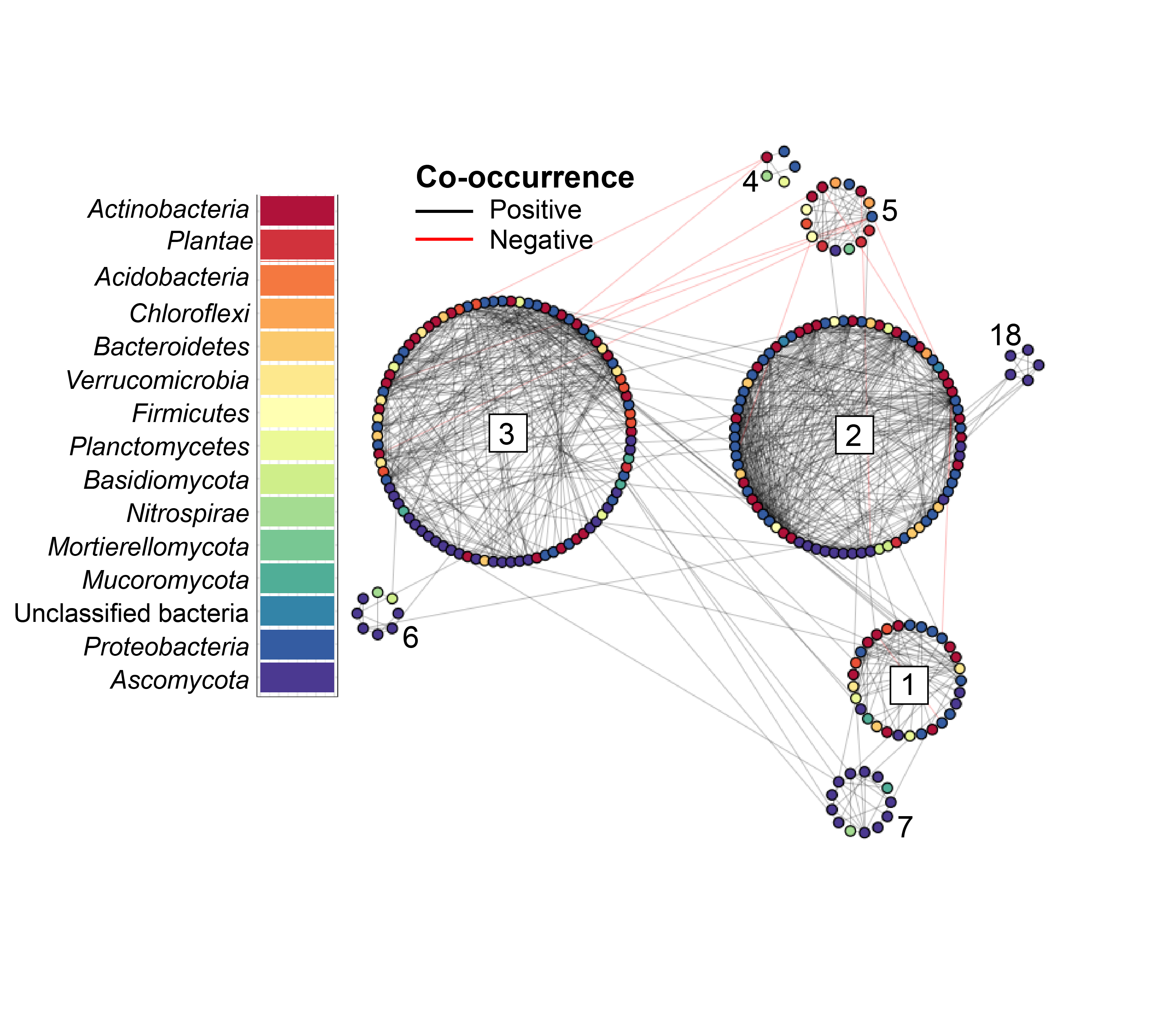

Supplemental Figure 27. Co-occurrence network [16S, ITS2, and plants - Organic Horizons], arranged by random walk modules. Each node represents an OTU and is coloured by phylum. Lines between points indicate co-occurrences (black) or co-exclusion (red). Module IDs are indicated with numbers for reference.

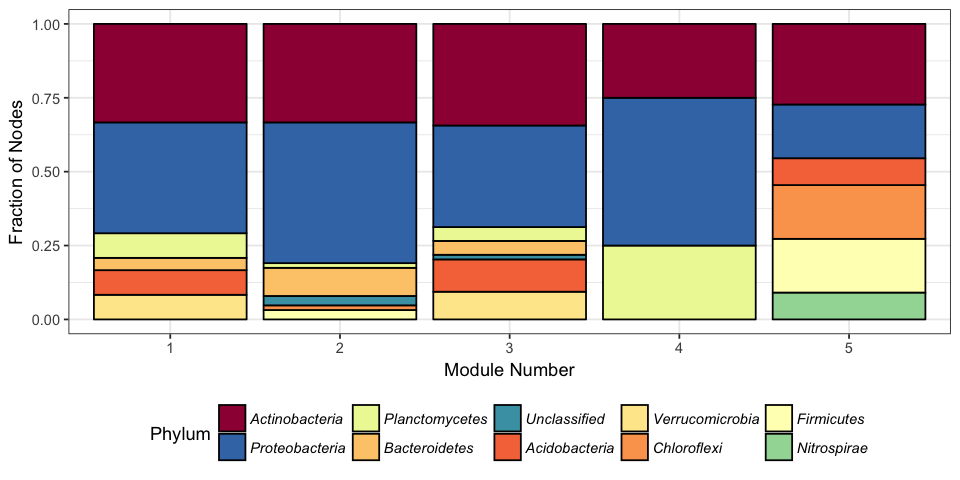

Supplemental Figure 28. Fraction of 16S OTUs in each module from different bacterial phyla in co-occurrence network [16S, ITS2, and plants - Organic Horizons].

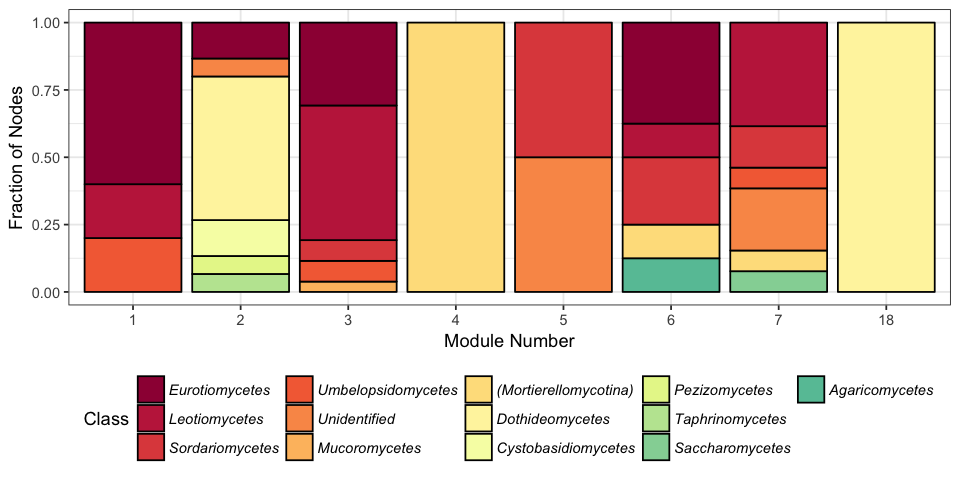

Supplemental Figure 29. Fraction of ITS2 OTUs in each module from different fungal classes in co-occurrence network [16S, ITS2, and plants - Organic Horizons].

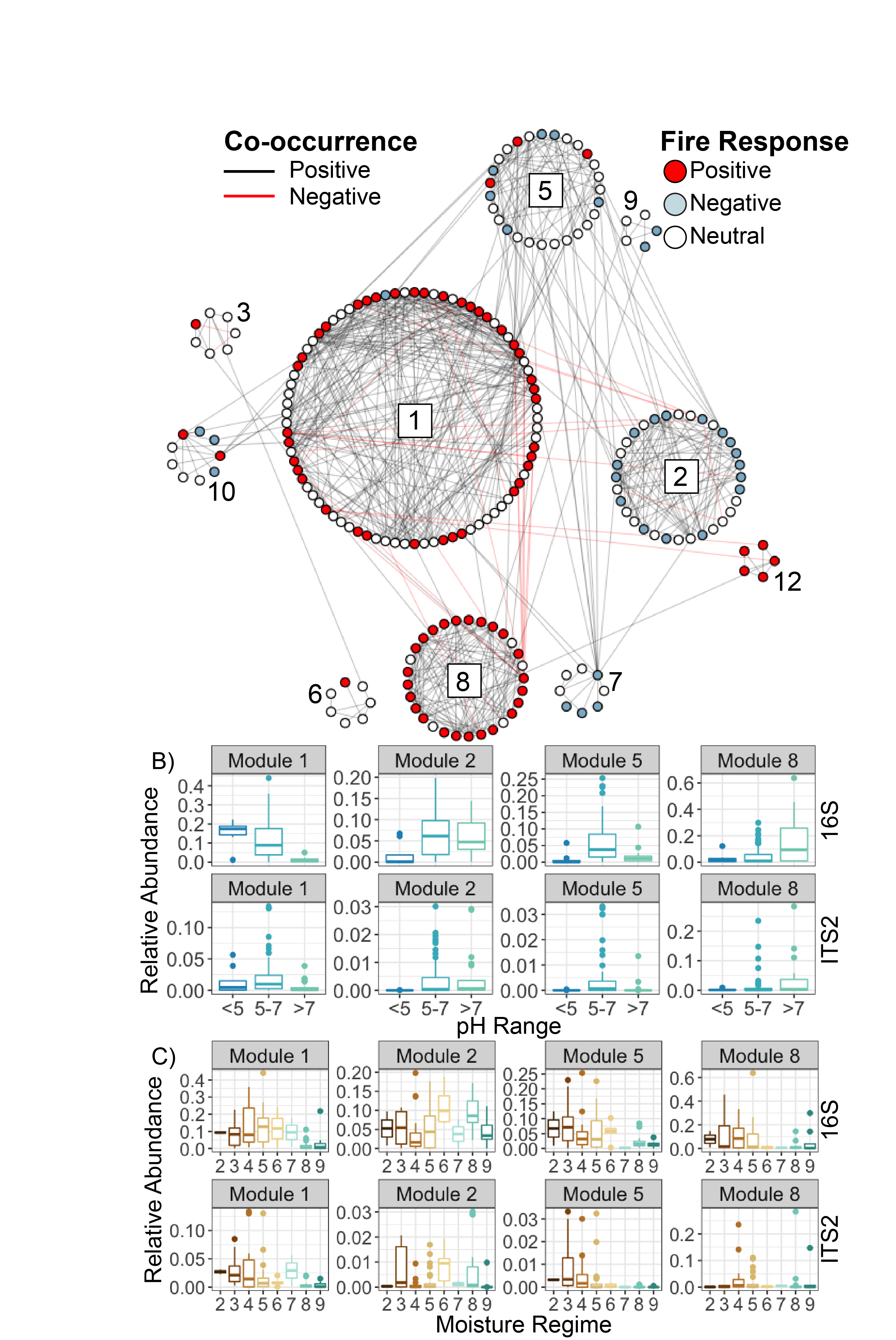

Supplemental Figure 30. A) Co-occurrence network [16S, ITS2, and plants - Mineral Horizons], arranged into greedy clustering-defined modules. Each point represents an OTU. Points are coloured by whether they were identified as being significantly more abundant in burned samples (red) and those significantly less abundant in burned samples (light blue) or no significant response (white). Lines between points indicate co-occurrences (black) or co-exclusion (red). Module IDs are indicated with numbers for reference. B) Module representation across moisture regimes: Fraction of total community represented by all bacterial (top, 16S) and fungal (bottom, ITS2) OTUs within selected modules, grouped by moisture regime, 2 being very dry, and 9 being very wet. C) Module representation across pH values: Fraction of total community represented by all bacterial (top, 16S) and fungal (bottom, ITS2) OTUs within selected modules grouped by pH range.

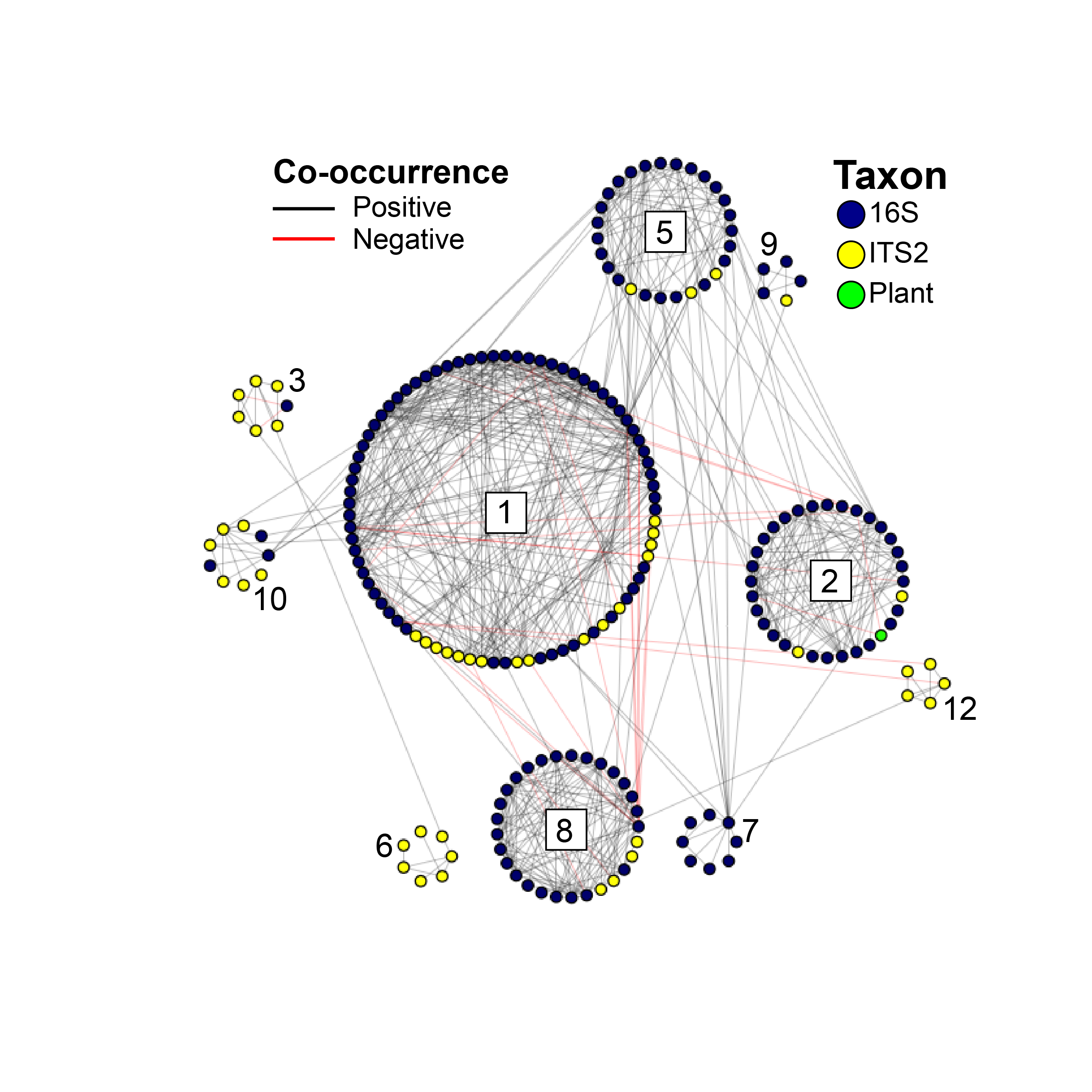

Supplemental Figure 31. Co-occurrence network [16S, ITS2, and plants - Mineral Horizons], arranged by random walk modules. Each point represents an OTU. Yellow=Fungi (ITS2), Navy=Bacteria and Archaea (16S), Green= Plants. Lines between points indicate co-occurrences (black) or co-exclusion (red). Module IDs are indicated with numbers for reference.

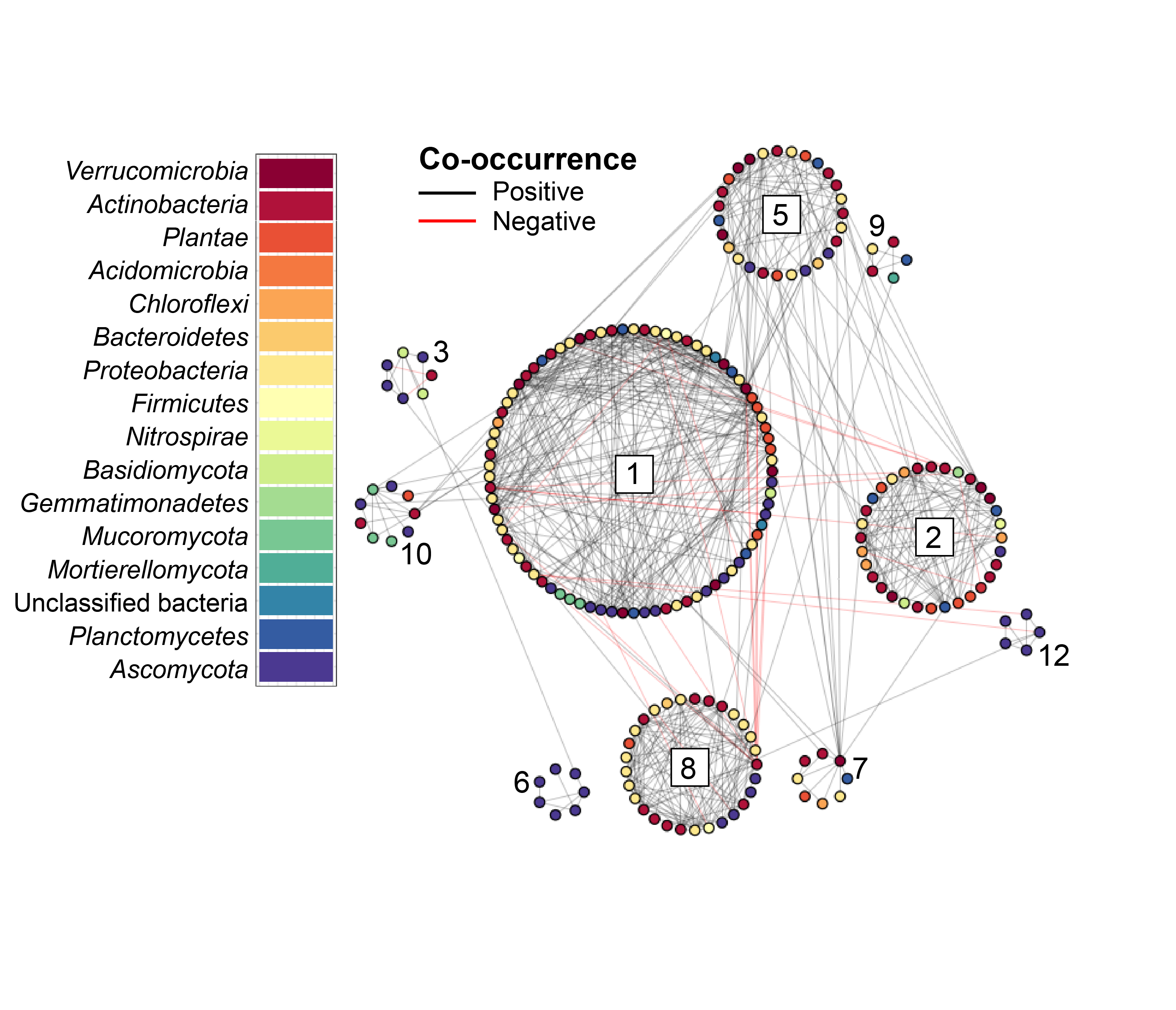

Supplemental Figure 32. Co-occurrence network [16S, ITS2, and plants - Mineral Horizons], arranged by random walk modules. Each node represents an OTU and is coloured by phylum. Lines between points indicate co-occurrences (black) or co-exclusion (red). Module IDs are indicated with numbers for reference.

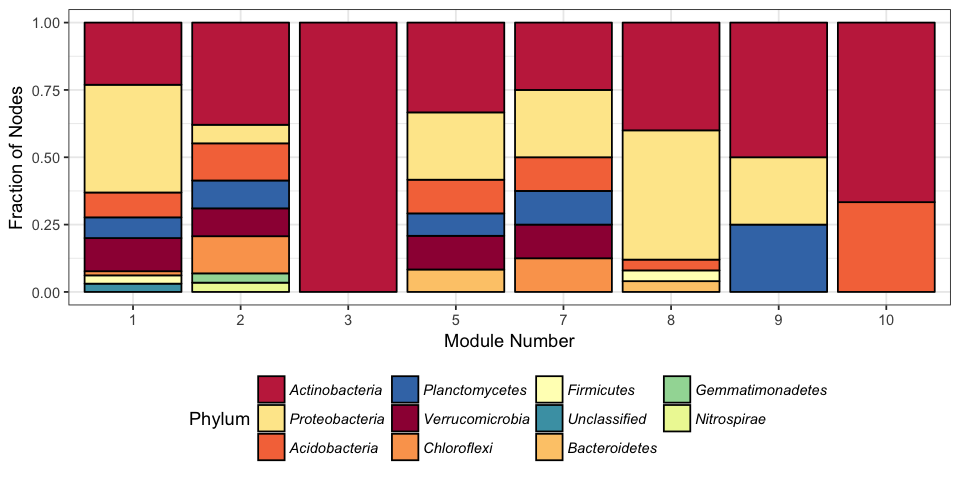

Supplemental Figure 33. Fraction of 16S OTUs in each module from different bacterial phyla in co-occurrence network [16S, ITS2, and plants - Mineral Horizons].

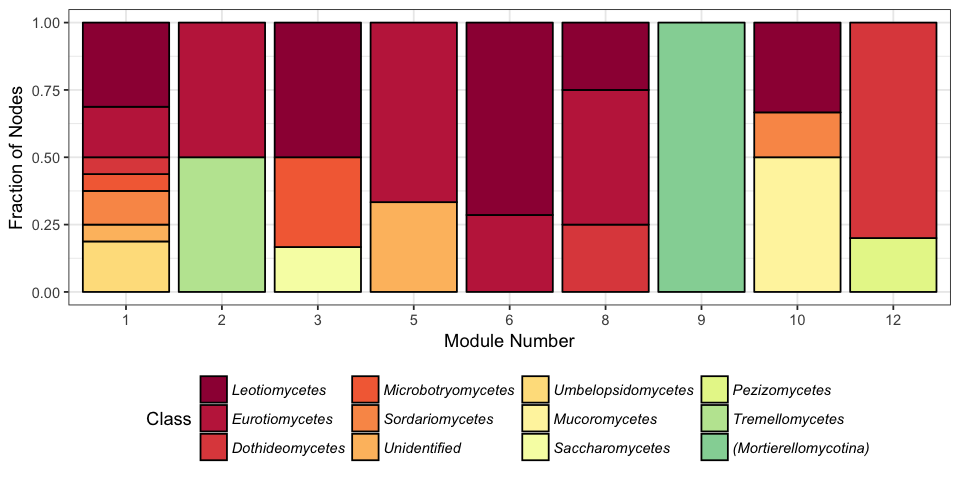

Supplemental Figure 34. Fraction of ITS2 OTUs in each module from different fungal classes in co-occurrence network [16S, ITS2, and plants - Mineral Horizons].

| Supplemental Table 1. Site characterization summary. Mean values reported with standard deviation in parentheses. Values reported only for burned sites. | | | | | |
| --- | --- | --- | --- | --- | --- |
|  | *Vegetation Community* | | | | |
| Vegetation Community | Black Spruce | Jack Pine | Mixedwood | Open Wetland | Treed Wetland |
| Total burned (unburned) | 6 (2) | 2 (15) | 15 (2) | 2 (11) | 11 (4) |
| Mean CBI | 2.1 (0.8) | 2.3 (0.7) | 1.6 (0.8) | 0.9 (0.4) | 2.2 (0.3) |
| Understory CBI | 1.7 (0.7) | 2.3 (0.6) | 1.5 (0.7) | 0.8 (0.3) | 1.8 (0.4) |
| Overstory CBI | 2.6 (1) | 2.3 (1) | 1.6 (0.9) | NA (NA) | 2.8 (0.4) |
| BSI | 3.5 (0.3) | 4.2 (0.7) | 3.6 (0.9) | 2.2 (0.4) | 3.1 (0.6) |
| CFSI | 4 (1.6) | 2.5 (2) | 1.6 (1.7) | 0.9 (1) | 3.2 (1.4) |
| RBR | 509.2 (123.1) | 397.9 (136.5) | 304 (153.7) | 112.6 (122.5) | 404.7 (97.1) |
| Exposed mineral soil (%) | 6.9 (9.1) | 41.4 (38) | 28.3 (39.8) | 0 (0) | 4.8 (6.9) |
| Duff depth (cm) | 6.6 (2.5) | 0.9 (1.4) | 1.9 (1.4) | 9.9 (0.4) | 8.8 (2.5) |
|  | *Overstory composition (stems ha^-1^)* | | | | |
| *Pinus banksiana* | 118 (289) | 1037 (1183) | 1735 (3460) | 10 (13) | 0 (0) |
| *Picea mariana* | 3946 (1083) | 452 (977) | 78 (164) | 32 (56) | 966 (992) |
| *Picea glauca* | 87 (212) | 9 (27) | 326 (691) | 6 (10) | 156 (270) |
| *Larix laricina* | 0 (0) | 1 (5) | 0 (0) | 27 (45) | 324 (371) |
| *Populus tremuloides* | 30 (72) | 119 (221) | 672 (1063) | 0 (0) | 17 (46) |
| *Populus balsamifera* | 0 (0) | 0 (0) | 10 (32) | 0 (0) | 0 (0) |
| *Betula papyrifera* | 81 (138) | 2 (7) | 8 (26) | 12 (38) | 0 (0) |

| Supplemental Table 2. Soil properties | | | | |
| --- | --- | --- | --- | --- |
| Property | Mean | Standard Deviation | Maximum | Minimum |
| *Organic horizon samples (n=59)* | | | | |
| pH | 6.2 | 1.0 | 7.7 | 3.2 |
| EC (mS cm^-1^) | 0.8 | 0.9 | 3.5 | 0.1 |
| Total C (%) | 33.3 | 15.0 | 52.8 | 3.4 |
| Total organic C (%) | 35.8 | 15.3 | 1.0 | 5.3 |
| Total N (%) | 1.2 | 0.8 | 2.7 | 0.02 |
| Total Ca (mg kg^-1^) | 28261.0 | 33753.1 | 211123.0 | 2922.8 |
| Total K (mg kg^-1^) | 746.9 | 506.5 | 2506.8 | 209.6 |
| Total Mg (mg kg^-1^) | 3094.4 | 4449.4 | 32701.7 | 423.4 |
| Total Na (mg kg^-1^) | 206.8 | 160.5 | 1020.1 | 88.5 |
| Total P (mg kg^-1^) | 798.2 | 590.5 | 4092.7 | 234.7 |
| Total S (mg kg^-1^) | 4100.3 | 5467.2 | 22357.7 | 253.2 |
| Total Al (mg kg^-1^) | 2413.7 | 4751.8 | 34457.3 | 210.8 |
| Total Fe (mg kg^-1^) | 3507.9 | 4287.3 | 19881.0 | 411.4 |
| Total Zn (mg kg^-1^) | 74.5 | 96.4 | 348.1 | 4.4 |
| Total Cu (mg kg^-1^) | 92.3 | 68.1 | 453.5 | 35.3 |
| Total Mn (mg kg^-1^) | 550.0 | 576.3 | 2509.7 | 18.6 |
| Total Mo (mg kg^-1^) | 118.3 | 314.0 | 2296.1 | 40.8 |
| O depth (cm) | 2.7 | 3.0 | 8.5 | 0.2 |
| *Mineral horizon samples (n=43)* | | | | |
| pH | 6.4 | 0.9 | 8.1 | 4.3 |
| EC (ms cm^-1^) | 0.2 | 0.4 | 2.4 | 0.0 |
| Total C (%) | 4.0 | 5.5 | 28.2 | 0.5 |
| Total organic C (%) | 4.5 | 6.0 | 1.0 | 0.5 |
| Total N (%) | 0.2 | 0.3 | 1.2 | 0.02 |
| Total S (mg kg^-1^) | 0.5 | 2.9 | 18.6 | 0.0 |
| Exchangeable Na (mg kg^-1^) | 51.3 | 10.6 | 90.7 | 38.5 |
| Exchangeable Mg (mg kg^-1^) | 254.2 | 256.0 | 985.0 | 29.2 |
| Exchangeable K (mg kg^-1^) | 150.3 | 71.3 | 392.4 | 84.5 |
| Exchangeable Ca (mg kg^-1^) | 3450.8 | 6420.0 | 39948.0 | 268.0 |
| Sand (%) | 58.8 | 29.2 | 95.0 | 22.0 |
| Silt (%) | 19.3 | 13.8 | 51.0 | 4.0 |
| Clay (%) | 7.2 | 7.7 | 33.0 | 1.0 |

| Supplemental Table 3. Primers used in this study (Burton *et al.*, 2008; de Groot *et al.*, 2013; Pellegrini *et al.*, 2017). Full Illumina PCR primers with barcodes listed in Supplemental Tables 4 and 5, attached as .csv files. | |
| --- | --- |
| **Primer** | **Sequence [**Illumina adaptor *Barcode* **Pad and linker** Primer] |
| *16S Illumina* |  |
| 515f | AATGATACGGCGACCACCGAGATCTACAC*XXXXXXXX***TATGGTAATTGT**GTGYCAGCMGCCGCGGTAA |
| 806r | CAAGCAGAAGACGGCATACGAGAT*XXXXXXXX***AGTCAGCCAGCC**GGACTACNVGGGTWTCTAAT |
| Read 1 seq | TATGGTAATTGTGTGYCAGCMGCCGCGGTAA |
| Read 2 seq | AGTCAGCCAGCCGGACTACNVGGGTWTCTAAT |
| Barcode seq | ATTAGAWACCCBNGTAGTCCGGCTGGCTGACT |
| *16S qPCR* |  |
| 515f | GTGYCAGCMGCCGCGGTAA |
| 806r | GGACTACNVGGGTWTCTAAT |
| *ITS2 Illumina* |  |
| ITS4 | AATGATACGGCGACCACCGAGATCTACAC*XXXXXXXX***TATGGTAATTAA**AGCCTCCGCTTATTGATATGCTTAART |
| 5.8S | CAAGCAGAAGACGGCATACGAGAT*XXXXXXXX***AGTCAGTCAGGG**AACTTTYRRCAAYGGATCWCT |
| Read 1 seq | TATGGTAATTAAAGCCTCCGCTTATTGATATGCTTAART |
| Read 2 seq | AGTCAGTCAGGGAACTTTYRRCAAYGGATCWCT |
| Barcode seq | AGWGATCCRTTGYYRAAAGTTCCCTGACTGACT |
| *ITS qPCR* |  |
| FR1 | AICCATTCAATCGGTAIT |
| FF390 | CGATAACGAACGAGACCT |

Supplemental Table 4. 16S Illumina PCR primers (available as .csv)

Supplemental Table 5. ITS2 Illumina PCR primers (available as .csv)

Supplemental Table 6. Fire-responsive 16S OTUs (available as .csv)

Supplemental Table 7. Fire-responsive ITS2 OTUs (available as .csv)

| Supplemental Table 8. Co-occurrence network properties for full networks | | | |
| --- | --- | --- | --- |
|  | *16S, ITS2, and plants* | | *16S and ITS2* |
|  | *Organic horizons* | *Mineral horizons* | *Organic and*  *mineral horizons* |
| Rho cutoff value used | 0.52 | 0.58 | 0.37 |
| Total nodes (taxa) | 289 | 279 | 591 |
| Total edges (co-occurrences) | 963 | 676 | 3454 |
| Average path length | 4.2 | 4.6 | 3.4 |
| Diameter (longest [shortest path between two nodes]) | 10 | 14 | 9 |
| Average degree (mean number of edges at nodes) | 6.7 | 4.8 | 11.7 |
| Edge density | 0.02 | 0.02 | 0.02 |
| Global clustering coefficient (all triangles) | 0.43 | 0.35 | 0.36 |
| Average clustering coefficient (each node) | 0.50 | 0.46 | 0.51 |
| Mean closeness centrality of nodes | 1.3x10^-4^ | 7.2x10^-5^ | 1.6x10^-4^ |
| R^2^ of power-law | 0.83 | 0.86 | 0.84 |
| Exponent (alpha) of power law | 1.3 | 1.4 | 1.2 |
| Modularity | 0.60 | 0.63 | 0.58 |

| Supplemental Table 9. Properties of co-occurrence network modules with more than 4 nodes for all three networks. Module ID is arbitrary and not comparable between networks. | | | | | | | |
| --- | --- | --- | --- | --- | --- | --- | --- |
|  | Nodes (n) | | | | Fraction Fire-Resp. | | Putative Saprotrophs |
| Module | Total | 16S | ITS2 | Plants | 16S & ITS2 | | ITS2 |
|  |  |  |  |  | (Positive) | (Negative) |  |
| *16S and ITS2 - Organic and Mineral Horizons* | | | | | | | |
| 1 | 83 | 63 | 20 | NA | 0.55 | 0.02 | 35% |
| 3 | 103 | 51 | 52 | NA | 0.12 | 0.27 | 42% |
| 4 | 135 | 92 | 43 | NA | 0.01 | 0.63 | 33% |
| 5 | 34 | 6 | 28 | NA | 0.12 | 0.00 | 29% |
| 8 | 5 | 0 | 5 | NA | 0.00 | 0.00 | 100% |
| 10 | 53 | 38 | 15 | NA | 0.04 | 0.28 | 40% |
| 11 | 111 | 71 | 40 | NA | 0.82 | 0.00 | 15% |
| 21 | 8 | 0 | 8 | NA | 0.00 | 0.12 | 13% |
| *Plants, 16S, and ITS2 - Organic Horizons* | | | | | | | |
| 1 | 29 | 24 | 5 | 0 | 0.00 | 0.41 | 40% |
| 2 | 78 | 63 | 15 | 0 | 0.85 | 0.03 | 27% |
| 3 | 91 | 64 | 26 | 1 | 0.48 | 0.06 | 38% |
| 4 | 5 | 4 | 1 | 0 | 0.00 | 0.80 | 0% |
| 5 | 15 | 11 | 2 | 2 | 0.00 | 0.46 | 0% |
| 6 | 8 | 0 | 8 | 0 | 0.50 | 0.00 | 50% |
| 7 | 13 | 0 | 13 | 0 | 0.00 | 0.31 | 23% |
| 18 | 5 | 0 | 5 | 0 | 0.80 | 0.00 | 0% |
| *Plants, 16S, and ITS2 - Mineral Horizons* | | | | | | | |
| 1 | 81 | 65 | 16 | 0 | 0.47 | 0.01 | 31% |
| 2 | 32 | 29 | 2 | 1 | 0.00 | 0.52 | 0% |
| 3 | 7 | 1 | 6 | 0 | 0.14 | 0.00 | 17% |
| 5 | 27 | 24 | 3 | 0 | 0.11 | 0.22 | 66% |
| 6 | 7 | 0 | 7 | 0 | 0.14 | 0.00 | 29% |
| 7 | 8 | 8 | 0 | 0 | 0.00 | 0.50 | NA |
| 8 | 29 | 25 | 4 | 0 | 0.82 | 0.00 | 25% |
| 9 | 5 | 4 | 1 | 0 | 0.00 | 0.40 | 100% |
| 10 | 9 | 3 | 6 | 0 | 0.22 | 0.33 | 50% |
| 12 | 5 | 0 | 5 | 0 | 1.00 | 0.00 | 0% |

Supplemental Table 10. Connector and hub taxa in co-occurrence network [16S and ITS2 - Organic and Mineral Horizons] (available as .csv)

Supplemental Table 11. Connector and hub taxa in co-occurrence network [Plants, 16S, and ITS2 - Organic Horizons] (available as .csv)

Supplemental Table 12. Connector and hub taxa in co-occurrence network [Plants, 16S, and ITS2 - Mineral Horizons] (available as .csv)

Supplemental Note 1. Full Methodological Details

*Study region*

We selected sites in the Northwest Territories and northern Alberta (Wood Buffalo National Park), Canada, and sampled them one year post-fire, in 2015 (Figure 1; Supplemental Table 1). The fires and the drivers of burn severity are described in detail in Whitman *et al*. (Whitman *et al.*, 2018b), while their effects on understory vegetation are described in detail in Whitman *et al*. (Whitman *et al.*, 2018a). The study region includes the boreal plain in the southwest, and the boreal shield in the northeast. The forests in the study region are dominated by jack pine (*Pinus banskiana* Lamb.), black spruce (*Picea mariana* (Mill.)), white spruce (*Picea glauca* (Moench) Voss), and trembling aspen (*Populus tremuloides* Michx.). Secondary species include eastern larch (*Larix laricina* (Du Roi) K. Koch), balsam poplar (*Populus balsamifera* L.), and paper birch (*Betula papyrifera* Marsh.). The study area also contains substantial peat-forming wetlands, covering about one third of the total area (Tarnocai *et al.*, 2002). We classified vegetation communities for each upland site as being jack pine-dominated, black spruce-dominated, or composed of a mix of coniferous and broadleaf trees (“mixedwood”). We classified vegetation communities for wetlands as open or treed, according to the Field Guild to Ecosites of Northern Alberta (Beckingham and Archibald, 1996). The study region has long, cold winters and short, hot summers, with mean annual temperatures between -4.3 °C and -1.8 °C and annual precipitation ranging from 300 to 360 mm (ESWG 1995, Wang et al. 2012). The fire regime of the study region includes infrequent stand-replacing fires every 40-350 years (Boulanger et al., 2012). These fires are typically small (<200 ha) (Stocks et al., 2002), but much of the burned area is contributed by a few large fires. The six large wildfires in this study ranged in size from ~14,000 to >700,000 ha. The soils in these regions are mostly classified as Typic Mesisols (32 sites), Orthic Gleysols (16 sites), or Orthic Gray Luvisols (8 sites) (Soil Landscapes of Canada map v.3.2). The sampled sites span a wide range of soil properties, with pH values ranging from 3.2 to 8.1, total C ranging from 0.5% (mineral horizon) to 52% (organic horizon), and a wide range of textures (Supplemental Table 2). Although the study area is within the discontinuous and sporadic permafrost zones of northern Canada (NRCan 1993), no sampled sites had frozen active layers in the top metre of soil.

*Site assessment methodologies*

Sites were selected and characterized as described in detail by Whitman *et al*. (2018a, 2018b). Briefly, field sites were located in areas > 100 m and < 2 km from roads and were a minimum of 103 m from each other (but on average 170 km apart). They were selected using stratified random sampling using burn severity as estimated from remote sensing. Additional sites were accessed opportunistically by helicopter and were selected to represent the local range of burn severity and vegetation communities, resulting in a total of 50 burned field sites. We selected an additional 12 control sites (not burned within the last 38 years before sampling, mean time since fire 95 years – “unburned”), chosen to reflect the range of vegetation communities sampled in the burned plots, for a total of 62 sites.

At each site we established a 30 x 30 m square plot with the vertical and horizontal orientations aligned with the cardinal directions, and 10 x 10 m subplots at the four corners. We measured post-fire organic horizon depth (up to 10 cm) at the inner corners of the 10 × 10 m subplots. Understory vegetation percent cover was assessed in five 1 x 1 m plots at the same four points as organic soil depth, and at the plot centre (Whitman *et al.*, 2018a). We assessed burn severity in the four subplots (Supplemental Table 1; described in detail in (Whitman *et al.*, 2018b)), using severity metrics of canopy fire severity index (CFSI; Kasischke et al., 2000), burn severity index (BSI; Loboda et al., 2013), and percent exposed mineral soil. We also assessed the composite burn index (CBI; understory, overstory, and mean; Key and Benson, 2006; Kasischke et al., 2008) in the entire 30 x 30 m plot area. We used the relativized burn ratio (RBR), to represent remotely sensed burned severity at each site. RBR was produced using multispectral Landsat 8 Operational Land Imager and Landsat 5 Thematic Mapper images (Landsat Level-1 imagery, courtesy of the USGS).

At each plot, we took soil cores (5.5 cm diameter, 8.5-13.5 cm depth) at three locations (centre, SW and NE subplots). Soil cores were gently extruded and separated into organic (O) horizons (where present) and mineral (M) horizons (where present in the top 13.5 cm of soil profile). The three samples were pooled by horizon at each site and mixed gently by hand in a bag. From these site-level samples, sub-samples were collected for microbial community analysis and stored in LifeGuard Soil Preservation solution (QIAGEN, Germantown, MD) in a 5 mL tube (Eppendorf, Hamburg, Germany). Tubes were kept as cold as possible while in the field (usually for less than 8h, but up to 2 days for remote sites) and then stored frozen. The remaining soil samples were air-dried and analyzed for a range of properties, including, pH and total carbon (C) (Supplemental Table 2).

*DNA extraction, amplification, and sequencing*

Duplicate DNA extractions were performed for each sample, with two blank extractions for every 24 samples (half of which were sequenced), using a DNEasy PowerLyzer PowerSoil DNA extraction kit (QIAGEN, Germantown, MD) following manufacturer’s instructions. Samples were lysed on a FastPrep-24 5G (MP Biomedicals, Santa Ana, CA) for 45 s at 6 m s^-1^. Extracted DNA was amplified in triplicate PCR, targeting the 16S rRNA gene v4 region with 515f and 806r primers (Walters *et al.*, 2015), and targeting the ITS2 gene region with 5.8S-Fun and ITS4-Fun primers (Taylor *et al.*, 2016) with barcodes and Illumina sequencing adapters added as per (Kozich *et al.*, 2013) (all primers in Supplemental Tables 3-5). PCR was performed with 12.5 μL Q5 Hot Start High-Fidelity 2X Master mix (New England BioLabs INC., Ipswich, MA), 1.25 μL 515f forward primer (10 μM), 1.25 μL 806r reverse primer (10 μM), 1 μL DNA extract, 1.25 µL BSA (20 mg mL^-1^) (VWR, Radnor, PA), and 7.75 μL PCR-grade water. The reactions took place on an Eppendorf Mastercycler nexus gradient (Hamburg, Germany) thermal cycler as follows: 98 °C for 2 minutes + (98 °C for 30 seconds + 58 °C for 15 seconds + 72 °C for 10 seconds) x 30 + 72 °C for 2 minutes and 4 °C hold. The PCR amplicon triplicates were pooled, purified and normalized using a SequalPrep Normalization Plate (96) Kit (ThermoFisher Scientific, Waltham, MA). Samples, including blanks, were pooled and library cleanup was performed using a Wizard SV Gel and PCR Clean-Up System A9282 (Promega, Madison, WI) according to manufacturer’s instructions except for the following two deviations (1) the SV Minicolumn incubation and centrifugation (steps 5.A.2-5.A.3) steps were repeated twice for each sample, and (2) nuclease-free water application was divided into 30 μL and 20 μL increments with the incubation step and centrifuge step after each addition (step 5.A.6). The pooled library was submitted to the UW Madison Biotechnology Center (UW-Madison, WI) for 2x250 paired end (PE) Illumina MiSeq sequencing for the 16S amplicons and 2x300 PE for the ITS2 amplicons. A brief note on the choice of ITS2 sequencing primers and method: from Taylor *et al*. (Taylor *et al.*, 2016), we expected some amplicons to be as long as at least 511 bp. Thus, in order to help obtain high-quality sequences of this length with MiSeq technology, we needed to use the Kozich *et al*. (Kozich *et al.*, 2013) primer design approach, where no sequencing effort is wasted on reading through PCR primers, and 2x300 PE sequencing, despite substantial quality drop-off toward the end of the reads. With a typical read merging success rate for ITS2 sequences of >90%, we were relatively satisfied with this approach.

*Sequence data processing and taxonomic assignments*

For 16S reads, we quality-filtered and trimmed (left trim 12 for both reads; truncation length 240 for forward reads, 190 for reverse reads, maxN=0, maxEE=2, truncQ=2, n=1e6), dereplicated, learned errors (4M reads, randomized), determined OTUs (pool=FALSE, omega A = 1x10^-40^, band size=16, minimum overlap=20), and removed chimeras (consensus method) using dada2 (Callahan *et al.*, 2016) as implemented in R. This quality control and OTU-picking process retained a median 71% of sequences across samples, resulting in a mean of 22,225 16S sequences per sample. For ITS2 reads, we first merged reads using PEAR (Zhang *et al.*, 2014), and then quality-filtered and trimmed (left trim 0 for both reads; truncation length 0 for both reads, maxN=0, maxEE=1, truncQ=2, n=1e6), dereplicated, learned errors (4M reads, randomized), determined OTUs (pool=FALSE, omega A = 1x10^-20^, band size=32, minimum overlap=20), and removed chimeras (consensus method) using dada2 (Callahan *et al.*, 2016). This quality control and OTU-picking process retained a median 94% of merged sequences across samples, resulting in a mean of 12,570 ITS2 sequences per sample. These sequence processing steps were performed on the UW-Madison Centre for High Throughput Computing cluster (Madison, WI). Upon initial analysis, we confirmed that the paired DNA extraction replicates were generally very similar to each other, compared to other samples. To avoid pseudo-replication, we combined the community composition data from paired extractions additively and proceeded with a single sequencing dataset for each soil sample. For 16S, this resulted in a mean of 49,789 sequences per soil sample, with a minimum of 8,064, and a maximum of 196,041. For ITS2, this resulted in a mean of 27,261 sequences per soil sample, with a minimum of 10,275, and a maximum of 89,370. Taxonomy was assigned to the 16S reads using a QIIME2 (Caporaso *et al.*, 2010) scikit-learn feature classifier trained on the 515f-806r region of the 99% ID OTUs from the Silva 119 database (Pruesse *et al.*, 2007) with default settings, removing 32 sequences that were not classified as Bacteria or Archaea. This resulted in a final count of 19,988 16S OTUs across the full dataset, 92% of which were classified to the phylum level and 56% of which were classified to the genus level. For the ITS2 reads, we first ran them through ITSx (Bengtsson-Palme *et al.*, 2013) to identify fungi and to remove plant sequences (97% of sequences were retained), and then assigned taxonomy using the UNITE species hypothesis 99% threshold database version 7.2, using the parallel_assign_taxonomy_uclust.py script in QIIME1 (Caporaso *et al.*, 2010) with default settings to the genus level. We also classified ITS2 taxonomic assignments using the FunGuild database (Nguyen *et al.*, 2016); 32% of OTUs received FunGuild assignments.

*Quantitative PCR*

To estimate the relative abundance of bacteria *vs.* fungi in a given sample, extracted DNA was amplified via quantitative PCR (qPCR) in triplicate, targeting the 16S rRNA gene v4 region with 515f and 806r primers (Carini *et al.*, 2016) and targeting the 18S gene region with FR1 and FF390 primers{ (Prévost-Bouré *et al.*) (all qPCR primers in Supplemental Table 3). qPCR reactions were manually performed in a UV-sterilized PCR Workstation. In Multiplate™ Low-Profile 96-Well Unskirted PCR Plates (cat# MLL9601, Bio-Rad Laboratories, Inc., Hercules, CA), reactions containing 20 µL total volumes had the following components: 10 µL SsoAdvanced Universal SYBR Green Supermix (Bio-Rad Laboratories, Inc., Hercules, CA), 1µL 515f or FR1 forward primer (10 µM), 1µL 806r or FF390 reverse primer (10 µM), 1 µL DNA extract (diluted 100^-1^ for 16S and 10^-1^ for 18S), 1 µL BSA (20 mg mL^-1^) (VWR, Radnor, PA), and 6 µL PCR-grade water. PCR plates were adhesively sealed with Microseal® 'B' Adhesive Seals (cat# MSB1001, Bio-Rad Laboratories, Inc., Hercules, CA). qPCR reaction conditions were optimized by testing the qPCR protocol with samples that contained a high concentration of DNA, a low concentration of DNA, a high concentration of DNA that had previously amplified poorly, and a low DNA concentration that had previously amplified successfully and amplifying on a gradient to find the optimal melting temperature. The reactions took place on an CFX96 Touch™ Real-Time PCR Detection System (Bio-Rad Laboratories, Inc., Hercules, CA), under the following cycling conditions: 98 °C for 3 min + (95 °C for 15 s + 60 °C for 30 s + plate read) x 45. A melt curve analysis was performed after thermocycling to ensure specificity: (65 °C for 5 s + 0.5 °C cycle^-1^ + plate read) x 60. Calibration standards and no-template controls (NTCs) were included on each plate in triplicate. Calibration standards were sourced from genomic DNA from a *Streptomyces sp.* isolate for 16S (5.58 x 10^9^ copies µL ^-1^) and a *Rhizoctonia solani* isolate for 18S (7.71 x 10^9^ copies µL ^-1^), which were aliquoted and stored in a -20 °C freezer in single-use quantities. Standard curves were serially-diluted to span 5.58 x 10^3^ – 5.58 x 10^7^ copies µL^-1^ for 16S and 7.71 x 10^2^ – 7.71 x 10^5^ copies µL ^-1^ for 18S. After amplification, raw data was downloaded from the thermocycler to be viewed in Bio-Rad CFX Manager 3.1 (Bio-Rad Laboratories, Inc., Hercules, CA) and exported as .xlsx files. Standard curve equations were generated by plotting the average Cq of each standard against the copy number in Microsoft Excel (version 16.12) and taking the logarithmic regression. Samples were re-amplified if Cq values were missing more than one value, variable within a sample, less than the highest standard, or greater than the NTC average (if NTC amplified), or if the R^2^-value of the calibration curve was less than 0.98. Raw Cq values and calibration curve results are available as Supplemental Data.

*Bioinformatics and statistics*

We worked primarily in Jupyter notebooks, with phyloseq (McMurdie and Holmes, 2013), ggplot (Wickham), and dplyr

(Wickham *et al.*) being instrumental in working with the data in R (R Core Team).

We compared community composition across samples using Bray-Curtis dissimilarities on Hellinger-transformed relative abundances (Legendre and Gallagher, 2001), which we represented using NMDS ordinations. We tested for significant effects of vegetation community, moisture regime, pH, carbon, texture (% sand), and burned/unburned using a permutational multivariate ANOVA (PERMANOVA; the adonis function in vegan (Oksanen *et al.*)). We expect that interactions between many of these parameters would be statistically significant, but including all of the possible terms would likely result in overparameterization of the model. Because the order of the terms in the PERMANOVA model affects the R^2^ of a given term, to compare the relative explanatory power of each component, we also compared the R^2^ of single-component models for each factor.

We predicted 16S rRNA gene copy numbers using the ribosomal RNA operon database (rrnDB) (Stoddard *et al.*, 2015). Briefly, we assigned taxonomy using the RDP database with a confidence cutoff of 0.8. Then, for all OTUs with a genus-level assignment, we used the mean copy number for that genus in the database as the predicted copy number. For OTUs without a genus-level assignment or that were not in the database, we used the mean copy number for all other taxa in this study. We then calculated the abundance-weighted mean predicted copy number for each sample using the approach of Nemergut *et al*.

(Nemergut *et al.*, 2016), where we divide OTU abundances by copy number, then multiply the adjusted relative abundances by the predicted copy number and sum the values for each sample. We tested for the relationship between weighted mean predicted copy number for each sample and burn severity (understory composite burn index) with a linear model.

To compare our findings with those of Holden *et al*. (Holden *et al.*, 2016), we calculated the mean understory CBI value for all sites at which each OTU within that phylum was present, and determined whether there were significant differences in these values between different fungal phyla, using an ANOVA and Tukey’s HSD for multiple comparison correction. We tested whether the ratio of fungal community Bray-Curtis dissimilarity to bacterial community Bray-Curtis dissimilarity was different across different severity class categories using ANOVA with Tukey’s HSD for multiple comparison correction. We tested whether the ratio of fungal community Bray-Curtis dissimilarity to bacterial community Bray-Curtis dissimilarity was different across different severity class categories using ANOVA with Tukey’s HSD for multiple comparison correction. We tested whether vegetation community dissimilarity was significantly correlated with bacterial or fungal community dissimilarity for all pairs of sites in mineral and in organic soil horizons using Mantel tests, with 999 permutations.

We wanted to examine the relative explanatory power of different burn severity metrics for predicting microbial community composition. To do this, we used a simple linear model, which controlled for parameters we expected to influence community composition – vegetation community, moisture regime (as a continuous variable), pH, texture (% sand), and total C – and then tested the inclusion of each severity metric, comparing the R^2^ values for the severity metric. Although we expect that significant interactions could exist between these parameters, we wanted to avoid over-fitting the model to the data. The severity metrics we tested included: Burned/Unburned, relativized burn ratio (RBR), Canopy Fire Severity Index (CFSI), Composite Burn Index (CBI), Understory CBI, Overstory CBI, Burn Severity Index (BSI), % exposed mineral soil, and mean duff depth.

We calculated the relationship between pH and log(16S copy number : 18S copy number) using a linear model in R. We estimated richness and its associated standard error in each sample using the *breakaway* function in R (Willis *et al.*, 2016).

We determined which OTUs were significantly enriched in burned plots (*vs.* unburned plots) using metagenomeSeq (Paulson *et al.*, 2013), after controlling for (including as variables) vegetation community (categorical variable), pH (continuous variable), and %C (continuous variable), resulting in an estimate of the log_2_-fold change in the abundance of each OTU in burned vs. unburned plots, across samples. For a small subset of OTUs, we investigated the relationship between their log(relative abundance) and burn severity (understory CBI as a continuous variable) using a linear model.

To determine which fungal and bacterial OTUs and understory vegetation co-occurred across samples, we used a network analysis approach, following Connor *et al*. (2017) to avoid false positives and establish conservative network cutoff parameters. We were interested in detecting co-occurrences between fungi, bacteria, and understory vegetation. Since we had two soil samples from most sites (organic and mineral horizons), but one vegetation community analysis, we did not see a clearly appropriate way to combine the community compositions from the two horizons for each site. Thus, we analyzed the combined datasets separately for all organic horizons and for all mineral horizons. In addition, we considered the full fungal and bacterial datasets from both horizons, without the plants. This resulted in three separate networks: organic horizons with microbes and plants; mineral horizons with microbes and plants, organic and mineral horizons with microbes and no plants. For each network, first, we combined the relative abundance tables for fungi, bacteria (and plants), retaining only taxa present at a total of 0.005 relative abundance, when summing all samples in the dataset. We added an offset of +2 to the normalized bacterial abundances and an offset of +4 to the normalized fungal abundances so that the ranking of the different datasets were preserved but never overlapped, despite being integrated into the same dataset. We then added random noise to the relative abundances at levels below the smallest difference between any two non-zero taxa. This allows for the breaking of ties. We calculated Spearman correlation coefficients (rho) between all pairs of taxa. We repeated this procedure 1000 times. We then repeated this procedure 1000 times with matrices that were randomly filled with the same abundance values (the null dataset). We can then determine an appropriate rho threshold value by plotting the fraction of OTUs that remain in the largest component of the network if we exclude correlations below x rho values, ranging from 0 to 1. Choosing rho above where this metric drops off dramatically in the null dataset helps ensure that co-occurrences that are essentially “noise” are excluded. In our dataset, this value lay around 0.48. Connor *et al*. recommend choosing a threshold slightly above this transition, but not so high that the network is not mostly disconnected (Connor *et al.*, 2017). Thus, we proceeded with a rho=0.5 threshold. Next, we calculated a set of parameters that characterize the network and compared these to the values in the null dataset, calculating the clustering coefficient (mean 0.32) and the average path length (mean 4.2). The range of both of these values fell significantly outside of the same values for the null dataset – *i.e.*, the network is more connected than would be expected by random chance. After these tests, we determined a consensus network by adding random tie-breaking noise to the matrix and using rho=0.5 as the cutoff 2000 times. We then selected only the co-occurrences that occurred in 95% of the 2000 replications. This represented the consensus network. We determined standard network characterization metrics, such as modularity (using random walks – the “walktrap” algorithm (Pons and Latapy, 2005)), average path length and average degree, and plotted the network using igraph R package (Csardi et al., 2006). For each node, we also calculated within-module connectivity (Z_i_) of each node and among module connectivity (P_i_) (Guimera & Amaral 2005), which we used to identify nodes that were module hubs (highly connected within a module; Z_i_ > 2.5) or connectors (nodes that connect different modules; P_i_ > 0.62) (Olesen et al. 2007; Zhou et al. 2010; Deng et al. 2012; Shi et al., 2016).

While it may be possible to use co-occurrence networks to infer true ecological relationships between organisms (*e.g.*, competition, predation, or keystone species) (Freilich *et al.*, 2018), we are very hesitant to interpret our network in this way, as the sampling design clearly mixes many taxa that likely never would have perceived each other or even truly shared a common microhabitat. Thus, we limit our interpretations of the network to causes and implications of co-occurrences, not of ecological interactions between nodes (taxa).

Supplemental Note 2. Significant predictors of community composition

Our observation that soil bacterial communities are more strongly structured by pH (than C), while C is a stronger predictor (than pH) for soil fungal communities, is consistent with previous findings (Fierer and Jackson, 2006; Bahram *et al.*, 2018). In addition, our finding that open wetlands strongly structured soil microbial communities is not surprising – the dramatically different hydrology and associated biogeochemistry of open wetlands would be expected to select for very distinct microbial communities, adapted for waterlogged, low-oxygen, high-carbon environments. We were somewhat surprised that the effects of burning stood out so clearly in the NMDS ordinations (triangles *vs*. circles in Figure 2) and was a significant predictor of microbial community, given the extraordinarily wide range of soil properties and vegetation communities that the study sites spanned. Numerous other researchers have found significant changes in soil microbial community composition after fires (*e.g.*, (Smith *et al.*, 2008; Hamman *et al.*, 2007), but most of these studies were limited to a smaller range of sites – for example, the Hamman *et al*. study consisted mostly of the same vegetation type (ponderosa pine (*Pinus ponderosa* Dougl. ex. Laws.) / slimstem muhly (*Muhlenbergia filiformis* Vasey)) and the range of pH values was small (6.7±0.1 standard error in unburned sites vs. 6.5±0.1 in burned sites) compared to this study, although pH was still a significant predictor of community composition. Smith *et al*. considered sites that had similar vegetation to each other (*Picea glauca* (Moench) Voss, *Populus tremuloides* Michx. and *Populus balsamifera* L.) in boreal forest in northern Alberta. They also found pH was a strong control on bacterial community composition (although its range is not reported), but that burning had a strong significant effect on community composition. Taş *et al*. (Taş et al., 2014) also found that pH and C were significant predictors of soil 16S community composition in a burned black spruce boreal forest. Regardless of fire status, pH and C are well-known controls on soil microbial community composition, and those trends are also observed in this study.

Supplemental Note 3. Fires did not significantly affect richness or fungal:bacterial ratios one year post-fire

It has previously been observed that fires may reduce fungal biomass by a greater fraction than bacterial biomass, due to differential susceptibility / tolerance (Holden and Treseder, 2013; Pressler *et al.*, 2018). We did not find a significant trend in the relative abundance of bacteria *vs.* fungi with increasing burn severity (Supplemental Figure 11). Although fungi are generally thought to be more readily killed by fire’s heat than are bacteria (Vázquez *et al.*, 1993; Pietikäinen, 2000; Mabuhay *et al.*, 2006; Dumontet *et al.*, 1996; Bååth *et al.*, 1995), filamentous fungi may also be better able to explore a wider volume of soil than non-filamentous bacteria, accessing regions that are more suitable for growth, and thus be less affected by post-fire conditions. These effects could balance each other out, or, as discussed above, those effects may not have been captured due to the timing of sampling, or perhaps all fires resulted in conditions sufficiently severe as to kill substantial numbers of microbes. For example, van der Voort *et al*. (van der Voort *et al.*, 2016) note that numerous soil bacteria are killed by temperatures as low as 50-80°C. Additionally, there would be a wide range of responses within these extremely broad groups (kingdom Fungi *vs.* domains Bacteria and Archaea) (Peay *et al.*, 2009); thus, looking for patterns in 16S *vs.*18S copy numbers may be too simplistic. Other studies have also noted that fire-imposed pH shifts can result in an increased proportion of bacteria to fungi (Bissett and Parkinson, 1980). Although we were unable to measure pH before and after burns, we did find a significant positive relationship between the relative abundance of bacteria vs. fungi with increasing pH in uplands (Supplemental Figure 12), which is consistent with previous findings. However, we cannot demonstrate that these relationships are due to changes in pH from fire; rather, they may be more due to differences in the initial soil conditions. Other studies have reported changes in microbial diversity post-fire, sometimes decreasing (Reazin *et al.*, 2016), sometimes increasing (Sun *et al.*, 2015; 2016), and sometimes not changing (Oliver *et al.*, 2015). We did not detect significant differences in OTU richness with fires or with increasing burn severity for bacteria or fungi (Supplemental Figures 13 and 14). Given that we would most likely expect to see these differences shortly after the fire, it may further suggest that the immediate, short-term effects on richness of direct killing by fire may be largely recovered. Additionally, we should note that accurately estimating richness and appropriately characterizing its uncertainty is notoriously difficult to do for soil microbial communities (Willis *et al.*, 2016), and so our failure to detect differences does not demonstrate that fire did not affect richness –rather, that we were unable to detect it in this dataset.

Supplemental Note 4. Discussion of fire-responsive fungal genera and phyla in other studies

Some of the fungal taxa we identified as positive fire-responders, such as *Neurospora* and *Geopyxis* are well-known post-fire fungi. *Neurospora sp.* spores in the soil may be stimulated by the heat from fire, and often rapidly proliferate under the bark of fire-killed trees, possibly being facilitated by transport by microfauna (Jacobson *et al.*, 2004). Vrålstad *et al.* (1998) suggest the widespread prevalence of *Geopyxis carbonaria* ascocarps post-fire may be driven by its escape from its fire-killed pine host, which, combined with its adaptations to the post-fire environment (saprotrophy and tolerance to high pH and low moisture), make it a common sight at burned sites. The findings of Greene *et al*. ( 2017) support this hypothesis – they found that tree damage and major duff reduction were required for *G. carbonaria* to produce a high abundance of ascocarps after a fire, in a study of a fire in a pine-spruce forest of the Kootenay National Park in the Rocky Mountains of British Columbia, Canada.

There were some canonical fire-responders that we did not identify as fire-responders – in particular, *Morchella spp.* – the morels. There are numerous reasons this could be the case. First, although many pyrophilic fungi have x	mycorrhizal lifestyles, we did not design our sampling scheme specifically to target tree root-associated fungi. Second, we only took three soil cores at each site. With high spatial variability in fungi, we may have missed some of taxa that were distributed more patchily. For example, *Morchella* made up as much as 2% of the soil fungal community in one sample, but were undetected in 64 samples. Third, the post-fire response may have already occurred for some of the common pyrophiles – Greene et al. (2017) found that the post-fire response of *Morchella* was still high one year post-fire, but dropped off dramatically in subsequent years.

Unlike Holden *et al*. (2016), for the fungal communities, we did not observe significant differences in mean CBI of sites at which OTUs were detected for *Ascomycota vs.* *Basididomycota* in uplands (Supplemental Figure 7A), but we did observe that the mean CBI for *Chytridiomycota* was significantly higher than all other phyla except *Glomeromycota* and *Mucoromycota*, while *Mucoromycota* was significantly higher than *Ascomycota, Mortierellomycota, and Rozellomycota*. Some *Chytridiomycota* have previously been noted to recover readily from heating as high as 90°C, which could provide a possible explanation for their increased relative prevalence at higher-severity burn sites (Gleason *et al.*, 2004). *Chytridiomycota* have also been noted as being dominant members of periglacial soil fungal communities (Freeman *et al.*, 2009). The soil conditions after very severe fires, where the O horizon is completely lost, might somewhat resemble that of plant-free high-elevation soils described by Freeman *et al*. (2009). We were concerned that this trend might be driven largely by moisture status, since the CBI mirrors the moisture regime (Supplemental Figure 7B). However, when we stratified the mean CBI calculations by moisture regime, the trend of *Chytridiomycota* being associated with high-CBI sites persisted (Supplemental Figure 8). Still, the phylum only makes up a small fraction of the total fungal community at burned sites, so the broader ecological importance of this effect should be considered circumspectly.

Supplemental Note 5. Discussion of fire-responsive bacterial genera in other studies

Although we did not test for significant differences in community composition at the phylum level, we did observe similar trends to previous studies - burned plots tended to have more *Actinobacteria*, *Bacteroidetes*, and *Firmicutes*, as well as *Betaproteobacteria* (Supplemental Figures 1 and 2) (Mikita-Barbato *et al*., 2014; Cobo-Diaz *et al*. 2015; Weber *et al*. 2014; Smith *et al*. 2008).

One of abundant fire-responsive bacteria was a *Blastococcus sp.* OTU, which was not detected at burned sites (Figure 4B). Cobo-Diaz *et al*. (2015) also found that *Blastococcus* was one of four genera that were enriched post-fire in a holm oak forest in the Sierra Nevada Natural and National Park in Spain. The same study also found *Bacillus* genus was enriched post-fire - we also found two *Bacillus* OTUs that were enriched with fire in this study (Supplemental Table 6). Using DGGE and targeted sequencing, Mikita-Barbato et al. (Mikita-Barbato *et al.*, 2015) found *Mycobacterium sp.* OTUs that were found in severely burned but not in the unburned comparison plots, as well as a *Mycobacterium sp.* OTU that was found in the unburned comparison plots, but not in the severely burned plots. Similarly, we identified two *Mycobacterium sp.* OTUs that were positive responders to fire, as well as five that were negative responders to fire. Weber et al. also found *Devosia*, *Pedobacter*, and *Adhaeribacter* genera were associated with both of their burned soils (mixed conifer and ponderosa pine), of which we found one, two and two positive burn-responding OTUs, respectively. In the ponderosa pine only, they also found *Sphingomonas* to be associated with burned soils, for which we identified two positively and one negatively responding OTUs, and *Massilia*, for which we identified three positively responding OTUs, including the most abundant OTU. In contrast, they found that *Acidobacteria* Group 4 was associated with burned ponderosa pine sites, while we only identified Group 4- associated OTUs as negative responders (all *Blastocatellaceae*). We also found *Acidobacteria* Subgroups 3 and 7 was negatively associated with burns (1 OTU and 2 OTUs, respectively) as was *Bradyrhizobium* (1 OTU). There were some taxa that Weber et al. (2014) identified as being fire-associated or not fire-associated, for which we observed opposite responses, but we note that the associations identified in the other paper were broadly qualitative, and might not have been identified as being statistically significant had the same analyses as this study been applied. In a fire chronosequence study in a boreal subarctic coniferous forest in northeastern Finland, Sun et al. found *Rhizobium* was only detectable at their 2-year post-burn site (vs. 60 or 152 years post-fire); we also identified one *Rhizobium* OTU as being associated with burned sites. In addition, the *Sphingomonas* genus was significantly more abundant in the 2-year vs 152-year post-fire sites; of this genus, we identified 2 positively-responding and 1 negatively-responding OTUs. Guo et al. also identified *Sphingomonas* as a potential PAH user in a PAH-contaminated soil study using 16S sequencing of DGGE bands (as well as *Massilia*) (Guo *et al.*, 2017).

Supplemental Note 6. *Calyptrozyma* as putative nutrient-responsive positive fire-responder

There were three OTUs identified as belonging to the genus *Calyptrozyma* that were also positive fire-responders. Searches on the Web of Science for Calyptrozyma+(soil or fire or boreal or pine or spruce) all yielded no results, and the taxon did not receive classification within the FUNGuild database (Nguyen *et al.*, 2016). However, OTU sq9 was a 100% match for OTUs found in soil associated with *Pinus radiata* in New Zealand (KX222617) (Addison *et al.*, 2018). In addition, it is 99% identical to an OTU that was detected in soils in an Alaskan black spruce forest, sampled four years after a moderately severe burn (HM164559; (Bent *et al.*, 2011)). However, the authors of that study make no specific observations about that taxon in the associated paper. It is also a 100% ID match to an OTU (GQ892250.1) identified as being significantly enriched after 5 years of N fertilization and 8 years after a fire in an Alaskan black spruce forest (Allison *et al.*, 2010). Thus, one possible interpretation of our findings could be that the positive fire response of *Calyptrozyma* observed here is due to increased post-fire nutrient availability. The relatively sparse data available about this abundant fire-responsive taxon suggest it could be an interesting target for future study.

Supplemental Note 7. Fire-responder similarity to PAH-degrading organisms

In addition to the possible ability to grow quickly post-fire, many of the most abundant fire-responsive taxa have 16S V4 regions that are genetically very similar to organisms that have been identified as PAH-degraders (See Supplemental Note 2 for details). For example, Guo *et al*. also identified two *Aeromicrobium* OTUs as being potential PAH users (Guo *et al.*, 2017), and alkane hydroxylase genes similar to those of *Aeromicrobium* were found to be abundant in the rhizoplane of grasses planted in a petroleum-contaminated soil (Tsuboi *et al.*, 2015). The fire-responder *Massilia sp.* OTU was a 99% ID match to an OTU (JF417747.1) included a database of coal seam-associated microbiota (Vick *et al.*, 2018) and a 100% match to two other *Massilia* *sp*. sequences deposited by the authors (MF670530 and MF628354). In addition, the OTU is a 100% ID match to a *Massilia alkalitolerans* (previously *Naxibacter alkalitolerans* (Xu *et al.*, 2005); KY010279) deposited under the title “Isolation of several PAH-degrading bacteria from petroleum contaminated soil samples”, but, unfortunately, we were not able to identify an associated published paper. Furthermore, the OTU is a 97% (KF573748; (Wang *et al.*, 2016)) and 98% (JX270637.1; (Liu *et al.*, 2014)) ID match for two different *Massilia* isolates identified as phenanthrene-degraders. The *Burkholderia-Paraburkholderia sp*. (OTU sq18) fire-responder is a 100% ID match for a *Burkholderia* that was isolated from a TNT-contaminated soil and identified as being able to degrade 2,4,6-trinitrotoluene (TNT) (Thijs *et al.*, 2018).

Supplemental Note 8. Co-occurrence network and ericoid fungi / ericaceous plants

One way that we had expected to verify the quality of our network was to look for expected co-occurrences between ericaceous plants and ericoid fungi. However, because of the stringency of our network development, only three plant taxa – *Salix*, *Carex*, and *Geranium* – were retained in any of the final networks. Understory plants were included in the final network in similar proportions as for bacteria and fungi (1-2% of all taxa), but the exclusion of all ericaceous plants from the network made this test impossible.

Wickham H. ggplot2: Elegant Graphics for Data Analysis. http://ggplot2.org.

Wickham H, François R, Henry L, Müller K. dplyr: A Grammar of Data Manipulation. https://CRAN.R-project.org/package=dplyr.

Wilhelm RC, Cardenas E, Maas KR, Leung H, McNeil L, Berch S, *et al.* (2017). Biogeography and organic matter removal shape long-term effects of timber harvesting on forest soil microbial communities. *Nature Publishing Group* 1–18.

Willis A, Bunge J, Whitman T. (2016). Improved detection of changes in species richness in high diversity microbial communities. *Journal of the Royal Statistical Society: Series C (Applied Statistics)* **10**: 1496.

Xu P, Li W-J, Tang S-K, Zhang Y-Q, Chen G-Z, Chen H-H, *et al.* (2005). Naxibacter alkalitolerans gen. nov., sp. nov., a novel member of the family ‘Oxalobacteraceae’ isolated from China. *International Journal of Systematic and Evolutionary Microbiology* **55**: 1149–1153.

Zhang J, Kobert K, Flouri T, Stamatakis A. (2014). PEAR: a fast and accurate Illumina Paired-End reAd mergeR. *Bioinformatics* **30**: 614–620.
